# Cell-based and cell-free firefly luciferase complementation assay to quantify Human Immunodeficiency Virus type 1 Rev-Rev interaction

**DOI:** 10.1101/2021.09.13.460071

**Authors:** Tucker Hansen, Jodie Baris, Min Zhao, Richard Sutton

**Affiliations:** Department of Internal Medicine Section of Infectious Diseases Yale School of Medicine, New Haven, CT 06520 United States of America

## Abstract

Rev is an essential regulatory protein of Human Immunodeficiency Virus type 1 (HIV) that is found in the nucleus of infected cells. Rev multimerizes on the Rev-response element (RRE) of HIV RNA to facilitate the export of intron-containing HIV mRNAs from the nucleus to the cytoplasm, and, as such, HIV cannot replicate in the absence of Rev. We have developed cell-intact and cell-free assays based upon a robust firefly split-luciferase complementation system, both of which quantify Rev-Rev interaction. Using the cell-based system we show that additional Crm1 did not impact the interaction whereas excess Rev reduced it. Furthermore, when a series of mutant Revs were tested, there was a strong correlation between the results of the cell-based assay and the results of a functional Rev trans-complementation infectivity assay. Of interest, a camelid nanobody (NB) that was known to inhibit Rev function enhanced Rev-Rev interaction in the cell-based system. We observed a similar increase in Rev-Rev interaction in a cell-free system, when cell lysates expressing NLUC-Rev or CLUC-Rev were simply mixed. In the cell-free system Rev-Rev interaction occurred within minutes and was inhibited by excess Rev. The levels of interaction between the mutant Revs tested varied by mutant type. Treatment of Rev lysates with RNAse minimally reduced the degree of interaction whereas addition of HIV RRE RNA enhanced the interaction. Purified GST-Rev protein inhibited the interaction. The Z-factor (Z’) for the cell-free system was ~0.85 when tested in 96-well format, and anti-Rev NB enhanced the interaction in the cell-free system. Thus, we have developed both cell-intact and cell-free systems that can reliably, rapidly, and reproducibly quantify Rev-Rev interaction. These assays, particularly the cell-free one, may be useful in screening and identifying compounds that inhibit Rev function on a high throughput basis.

## Introduction

Human immunodeficiency virus type 1 (HIV-1) affects more than 38 million individuals world-wide, and an estimated 1.7 million individuals acquired the virus in 2019. The introduction of highly active anti-retroviral therapy (HAART) in the mid 1990s has greatly improved the prognosis of those living with the virus, and HIV infection, which was once invariably fatal, is now a chronic disease that can be managed by life-long HAART [1, 2]. There are currently dozens of FDA-approved medications for treating HIV+ individuals, and most individuals on combination therapy do remarkably well, with undetectable viral loads, increased CD4+ T cell counts, and near-normal life expectancy [2, 3]. There is a subset of individuals, however, who do poorly due to drug toxicity or tolerability/adherence issues or because drug resistance occurs with virologic escape mutants that can replicate to high levels and result in reduced CD4+ T cell counts [4–24]. Therefore, as with any other chronic disease process, for HIV there is always a need for improvement in current medications or for new, ‘first-in-class’ antivirals. A drug that interferes with Rev’s known mechanism of action would be one such first-in-class medication, since none of the current FDA-approved drugs possess anti-Rev activity.

Rev is an 18-kDa regulatory protein found in the HIV genome that is highly conserved among all viral isolates and clades [25, 26]. The *rev* gene is in a different reading frame than the viral gene *tat* but shares the same major intron, and the resulting protein, Rev, is encoded by a completely spliced HIV mRNA. After cytosolic translation Rev is imported into the nucleus, where interferes with the splicing pathway of HIV RNA transcripts [27]. Specifically, Rev multimerizes on the Rev-response Element (RRE), a ~260 base structured RNA located within the *tat/rev* intron, in order facilitate nuclear export of unspliced and partially spliced viral RNAs, including HIV genomic RNA [27–31]. Based upon biochemical analyses, 6-8 Rev proteins cooperatively bind to a single RRE [32–38], and a low-resolution structure has demonstrated a Rev dimer binding across a bent RRE [39]. The multimerization domain of Rev is separate from its RNA binding domain; the latter is known as the Arginine-Rich-Motif (ARM) [40]. There exist loss-of-function Rev mutants, used herein, that are able to bind RNA but are unable to multimerize [28].

Host factors Crm1 and Ran loaded with GTP form a complex with the Rev multimers within the nucleus in order to facilitate the export of viral mRNA through intact nuclear pores [41, 42]. Once the viral RNA has been exported to the cytosol, the complex dissociates, and its components are recycled back to the nucleus to export additional cargo. In the absence of Rev, essential HIV proteins encoded by ‘late’ intron-containing mRNAs, such as Gag, Pol, and Env, are not exported from the nucleus, and these RNAs are either confined to the nucleus and turnover or get spliced. There are no cellular homologs of Rev, and only rarely are intron-containing cellular RNAs exported to the cytosol via an analogous system [43, 44]. The absolute necessity of Rev in HIV replication makes it an obvious drug target, and the multimerization of Rev is a key aspect of its mechanism of action that could potentially be disrupted by a small molecule.

Firefly luciferase (FFLUC) is a 61-kDa monomeric bioluminescent enzyme from the *P. pyralis* firefly. The enzyme is a single polypeptide with two major domains, deemed the N-terminal domain and the C-terminal domain [45]. Importantly, when FFLUC is split into its two domains, neither the N-terminal domain nor the C-terminal domain is enzymatically active by itself [46]. The loss of activity can be explained by biochemical analysis, which suggests that the active site of FFLUC is situated between the two major domains [45]. In the presence of oxygen, ATP, and Mg^2+^ FFLUC oxidizes its D-luciferin substrate to produce an oxyluciferin product and light [47]. This reaction is at the center of a variety of biochemical assays that have been used for decades to quantify biological processes, including transcription, translation, post-transcriptional regulation, and protein-protein interaction. FFLUC’s dynamic range of many orders-of-magnitude coupled with a clear readout of activity has lent itself to high throughput screens to identify inhibitory compounds [48]. In the HIV field, FFLUC activity has been routinely used in viral pseudotyping assays with TZMbl target cells to measure the potency and breadth of anti-HIV broadly neutralizing antibodies, in addition to quantifying antibody-dependent cellular cytotoxicity or ADCC [49].

Because FFLUC is modular, more than 15 years ago a split-luciferase complementation assay (SLCA) was developed to assess protein interaction [50]. In this assay the N-terminal domain of FFLUC (NLUC) is fused to the C-terminus of gene A, and the C-terminal domain of FFLUC (CLUC) is fused to the N-terminus of gene B [50]. The enzymatic activity of FFLUC is restored, at least in part, when proteins A and B interact. In the initial report A=FRB and B=FKBP, and the interaction/enzymatic activity was dependent upon the addition of the small molecule rapamycin [50]. Here, we decided to test the utility of the SLCA in quantifying the ability of HIV Rev to interact with itself, where A=B=full length Rev (116 amino acids). We first tested this system under cell-intact conditions in which Rev-NLUC and CLUC-Rev were expressed in the same cell, but the system also worked under cell-free conditions, when individual Rev-NLUC and CLUC-Rev cell lysates were mixed. The cell-free system, in particular, is easily amenable to establishment of a high throughput to identify compounds that are capable of disrupting Rev-Rev interaction and hence Rev function.

## Results

### Construction and testing of the Rev-NLUC and CLUC-Rev fusions in the SLCA

We wished to determine whether the SLCA could be used to quantify Rev’s ability to interact with itself. Full-length Rev from HIV isolate NL4-3 was PCR amplified, sequence-confirmed, and inserted in frame, separately replacing FRB in FRB-NLUC and FKBP in CLUC-FKBP. These plasmids were transiently transfected into 293T cells, and expression of each fusion was confirmed by RIPA lysis of the cells at 72 h, followed by SDS-PAGE and immunoblotting using anti-FFLUC antisera. The sizes of the Rev-NLUC and CLUC-Rev fusions were ~70 kDa and ~35 kDa, respectively, as expected, with probing for GAPDH serving as loading control (Figure 1a-d). We next transiently transfected into 293T cells the two Rev fusion constructs separately or together, along with the parental FRB-NLUC and CLUC-FKBP plasmids. For the latter, 5 h after DNA transfection we treated the cells with rapamycin, and transfected cells were lysed at 72 h for FFLUC assay. In the presence of both Rev-NLUC and CLUC-Rev there was >100 fold increase in RLU compared to each plasmid alone; this was similar to what was observed with transfection of FRB-NLUC and CLUC-FKBP plasmids in the presence of rapamycin (Figure 2a). This result suggests that we are observing Rev-Rev interaction in this system, as assessed by the increased RLU.

**Figure 1.**
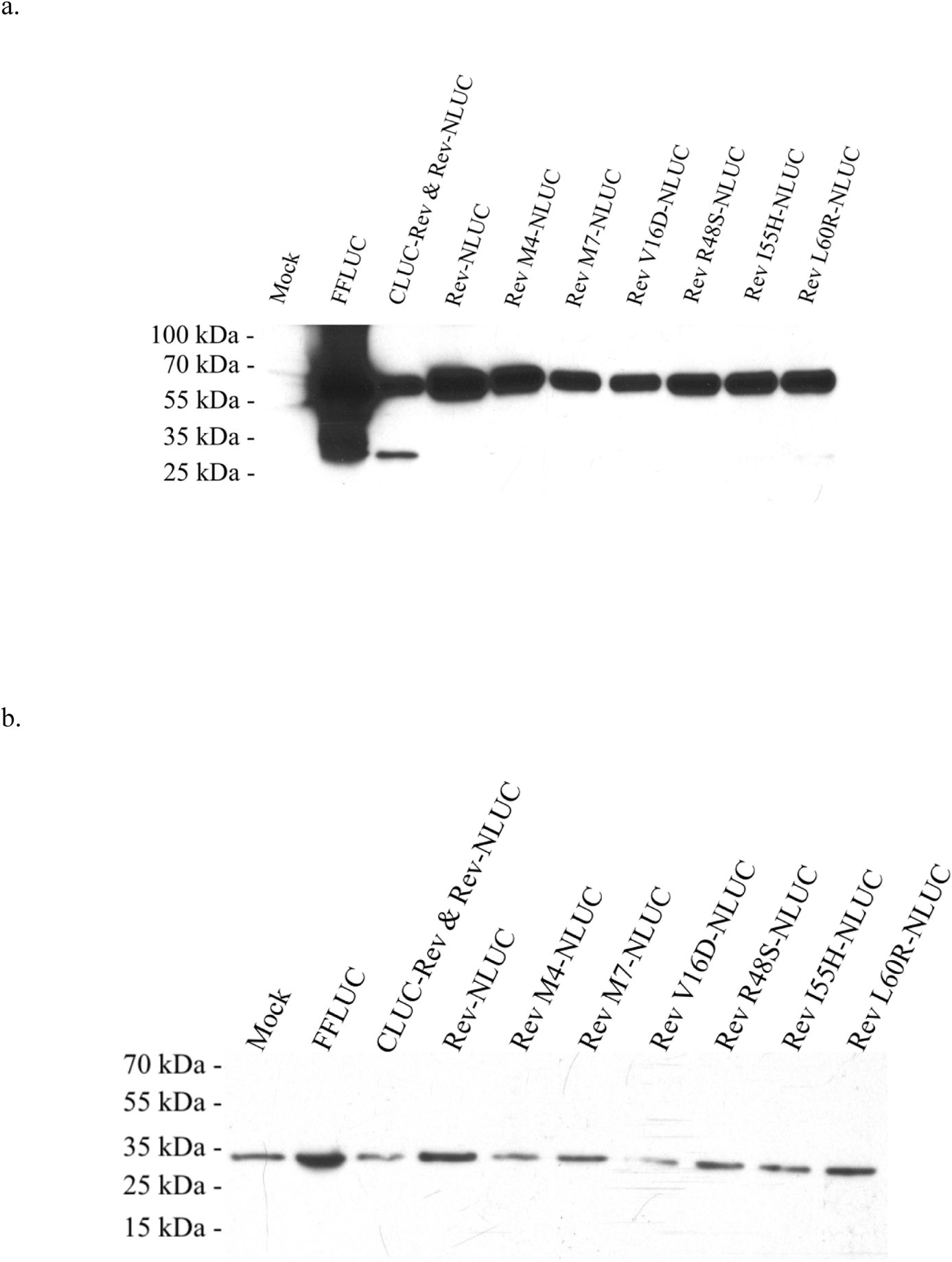
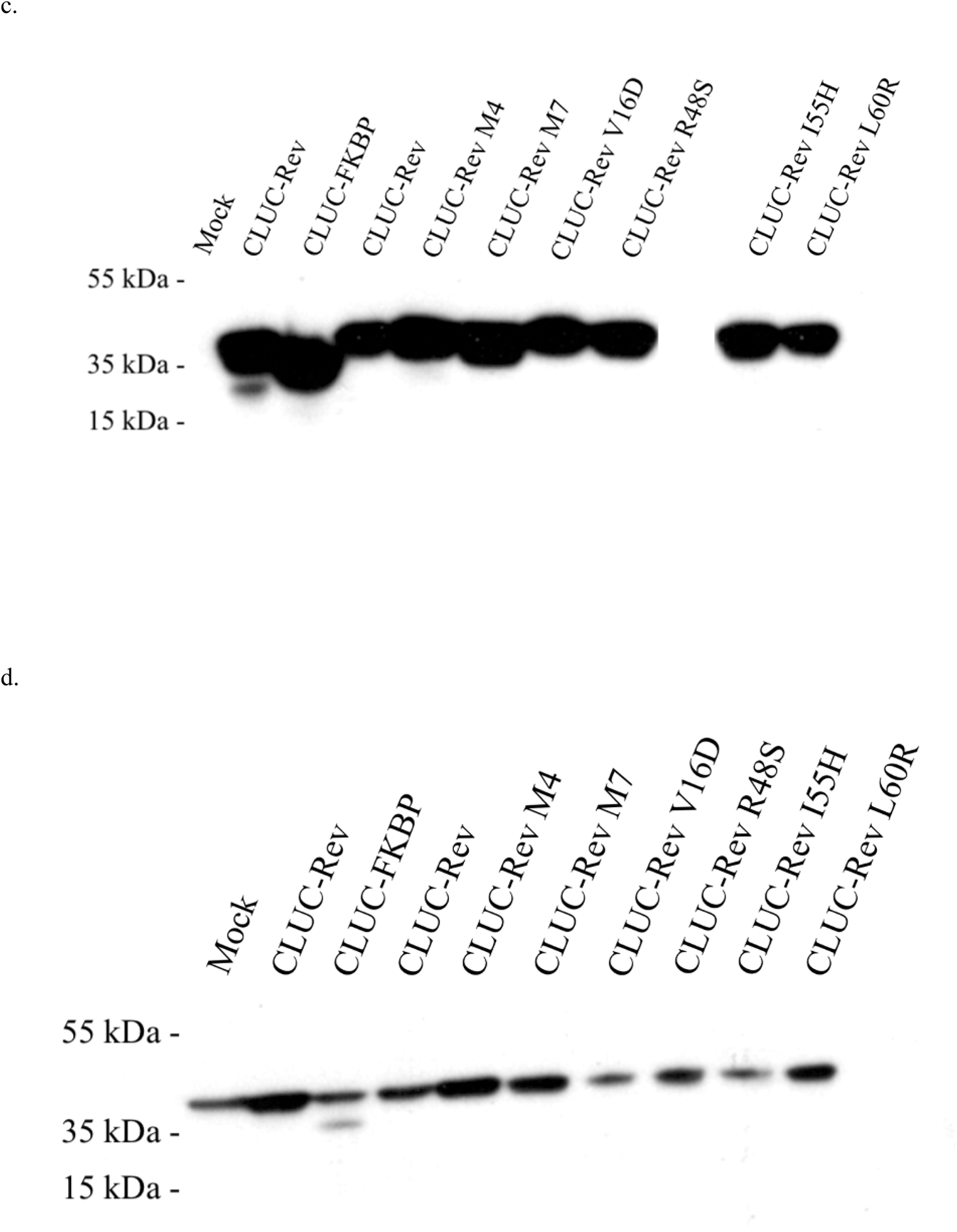
Expression of CLUC-Revs and Rev-NLUCs. 293Ts were transfected with indicated CMV-driven expression plasmids at top of each lane; mock refers to transfection in the absence of any plasmid. Protein ladder was run in parallel with sizes indicated. a) Immunoblot using anti-FFLUC antiserum. b) Corresponding immunoblot using anti-GAPDH antiserum. c) Immunoblot using Anti-FFLUC antiserum. Empty gel lane was removed between CLUC-Rev R48S and CLUC-Rev I55H. d) Corresponding immunoblot using anti-GAPDH antiserum.

**Figure 2.**
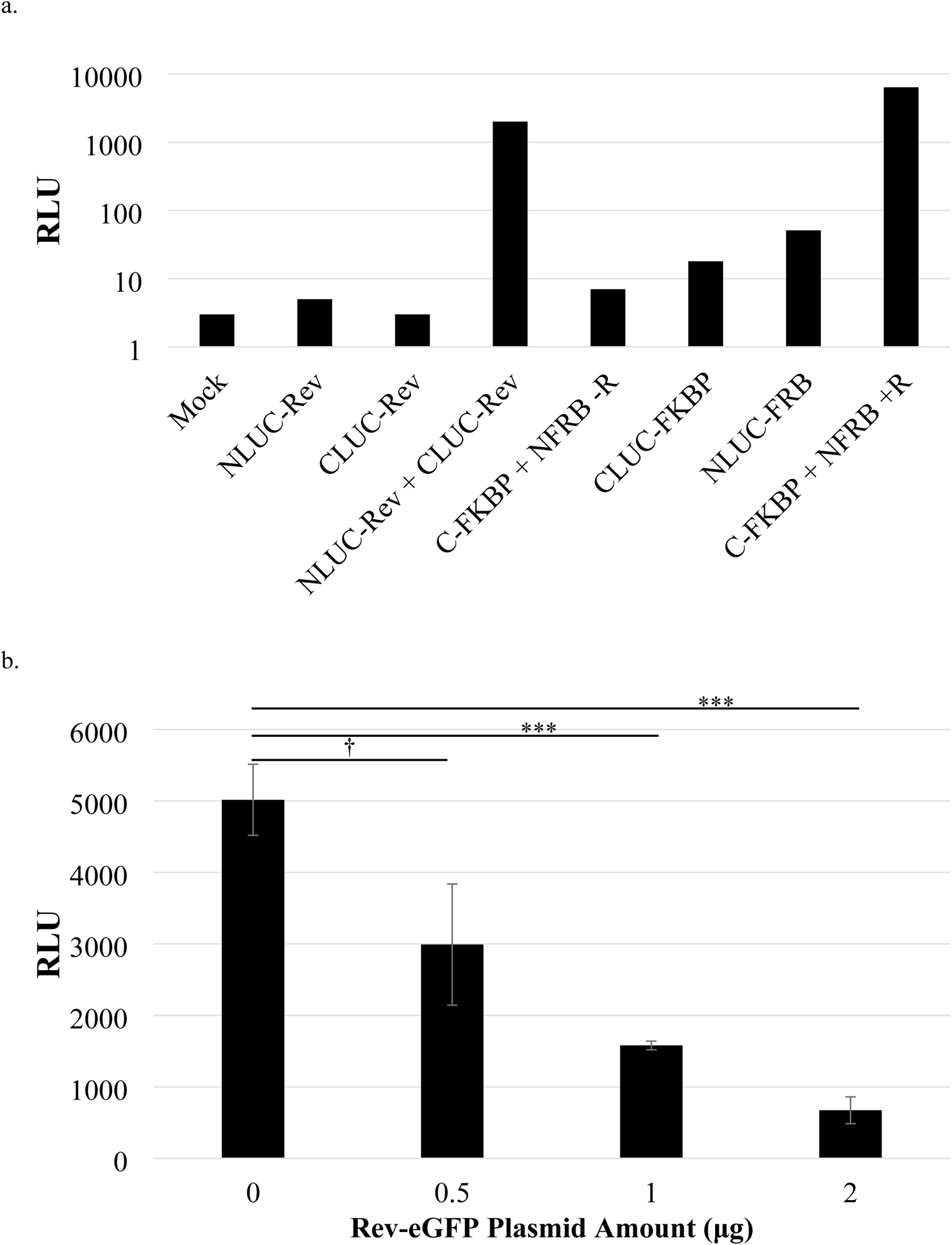
Co-transfection of Rev-NLUC and CLUC-Rev results in RLU. a) 293T cells were transfected with mock or indicated plasmids, in some cases with (+R) or without (-R) rapamycin. All plasmid constructs were CMV-driven. Cells were lysed at 72 h and assayed for luciferase activity. Note log scale for RLU. b) Both Rev-NLUC and CLUC-Rev plasmids were transfected into 293T cells in the presence of increasing amounts of Rev-eGFP fusion plasmid and RLU measured as in a). Values are mean ±SD. Student’s t-tests were performed to compare difference in means between each of the plasmid amounts and no plasmid control, correcting for multiple comparison testing. † denotes p >0.05, * p≤0.05, ** p≤0.01, *** p≤0.001, **** p≤0.0001.

If the observed interaction is truly due to affinity between Revs, we should be able to modulate it by competition with excess Rev. We transfected both Rev-NLUC and CLUC-Rev plasmids in the presence of increasing amounts of a Rev-eGFP fusion and observed an >80% reduction in RLU, which was significant at the higher amounts used (Figure 2b). This reduction cannot simply be attributed to transcriptional squelching since RLU was significantly lower when compared to transfection of empty vector (Figure 4c-d). For confirmation of the ~50 kDa Rev-eGFP protein expression, immunoblotting was performed using anti-Rev antisera (Figure 4a). Note these experiments were performed in the absence of any RRE RNA; thus, the observed interaction is RRE-independent.

**Figure 4.**
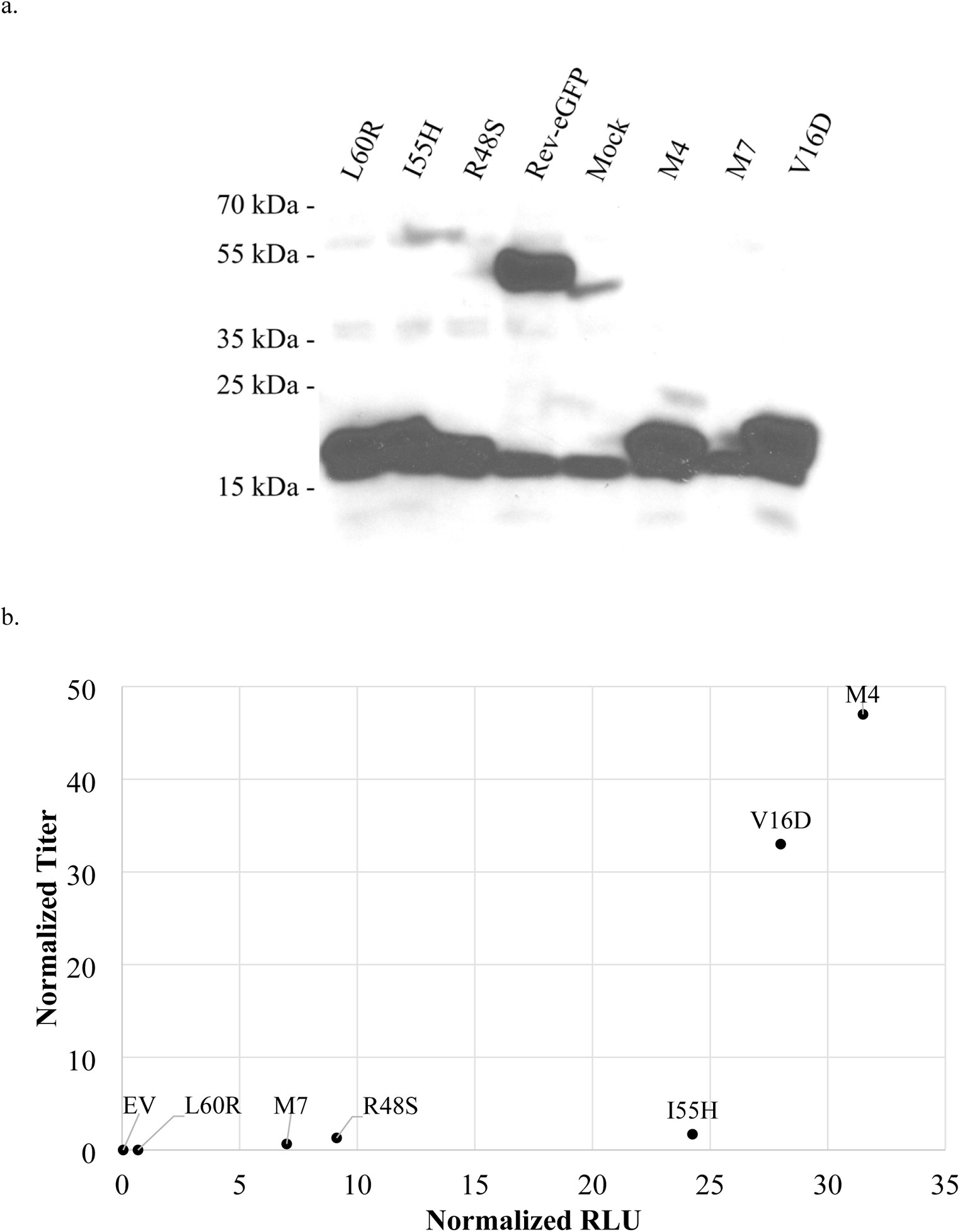
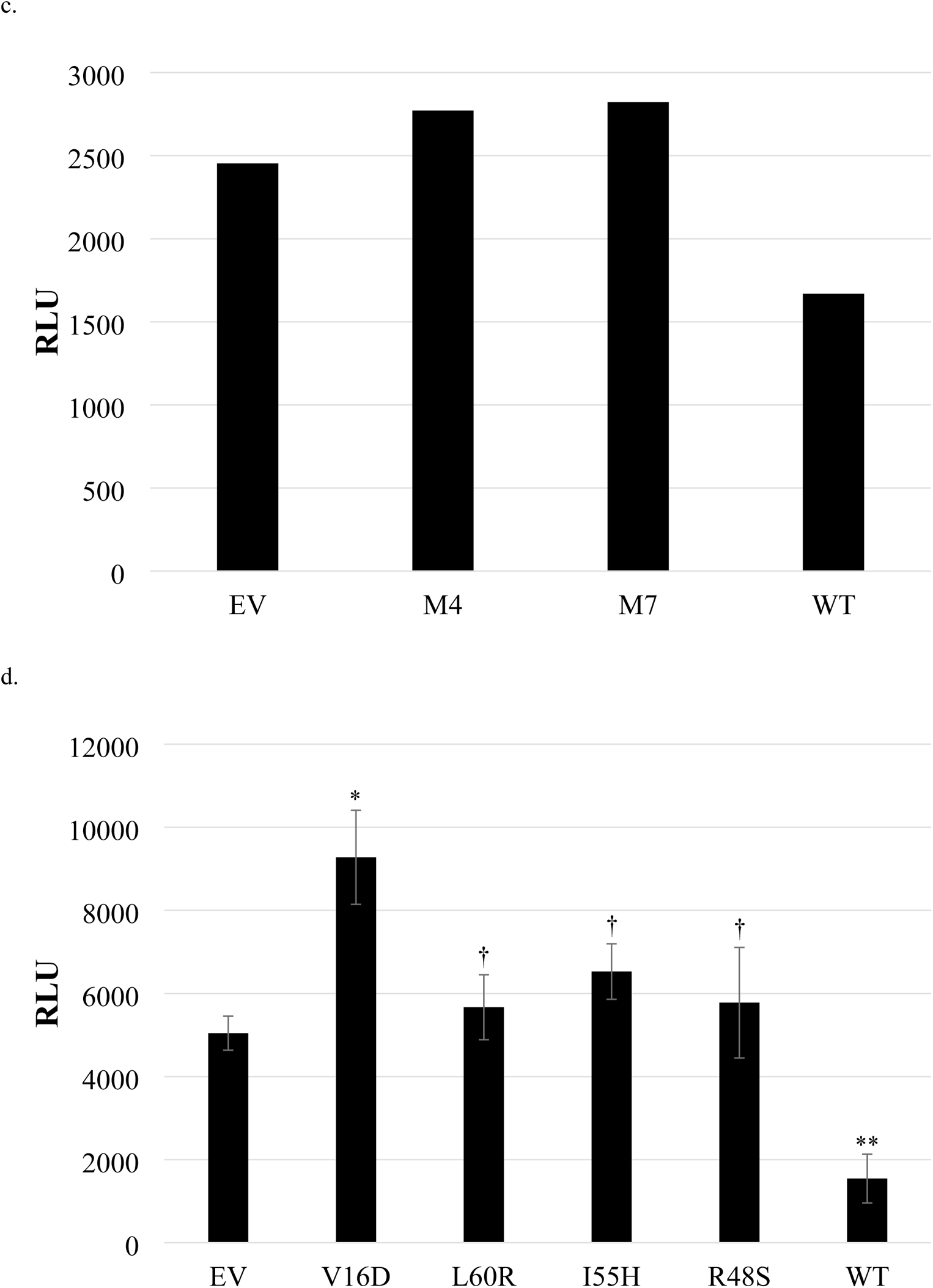
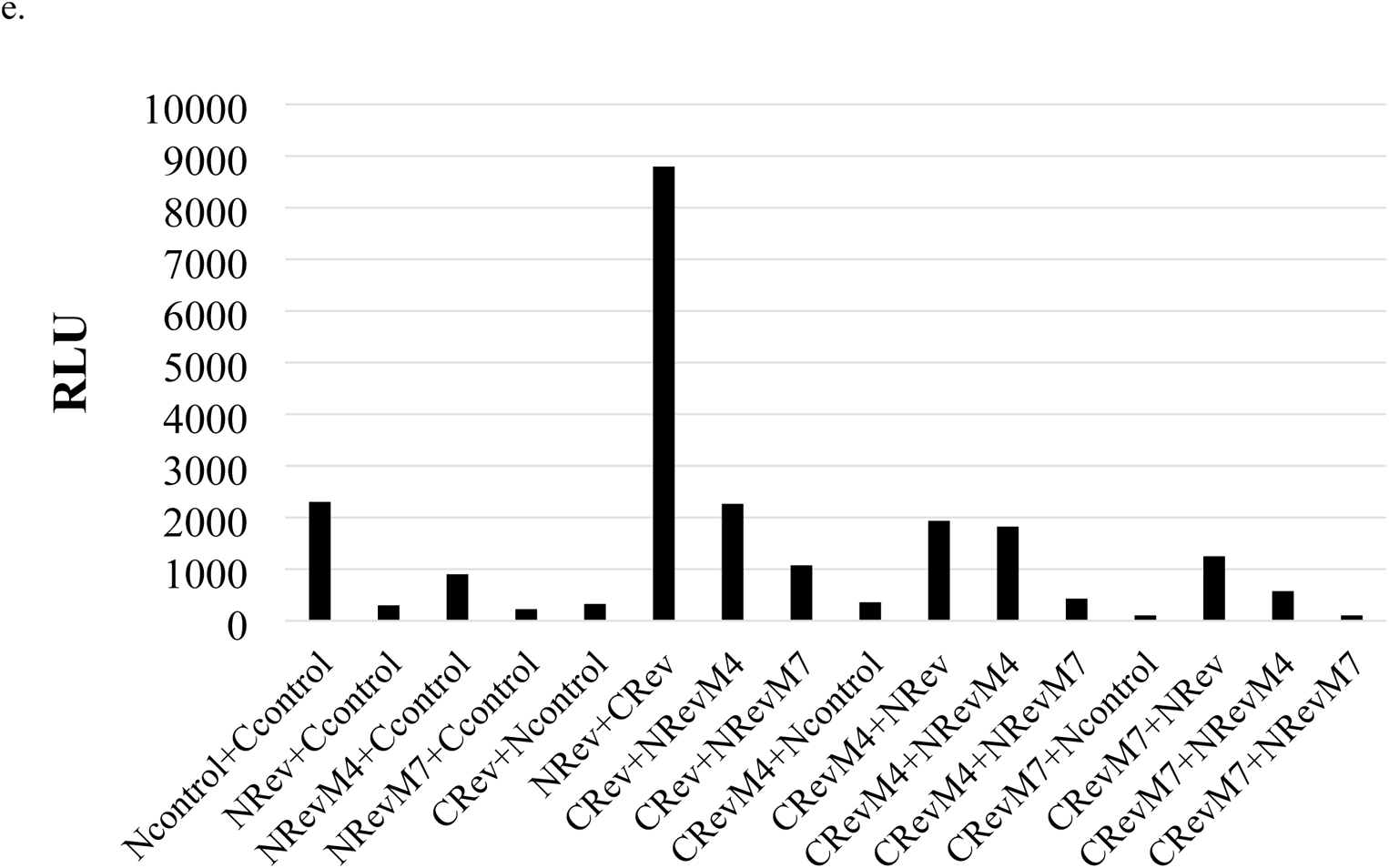
Rev trans-complementation infectivity assay and activity of mutant Revs. a) Immunoblot of mutant Rev plasmids indicated at top using anti-Rev antiserum. b) Correlation between viral titer and RLU. 293T cells were co-transfected with HIV-PVΔRev VSV G, a third generation HIV vector encoding *bsd^r^* gene, and, each, unfused Rev plasmid. Cell culture supernatant was harvested at 72 h and titered on 293T targets by serial dilution and enumerating blasticidin-resistant CFU. In parallel wt and mutant Rev-NLUC and CLUC-Rev expression plasmids were co-transfected into 293Ts and at 72 h RLU measured. Pearson r=0.81. From left to right: EV; L60R; M7=SIS→I, aa 54-56; R48S; I55H; V16D; M4=YSN→DDL, aa 23,25,26. Not shown is wt Rev (set at 100 on both axes). c) 293Ts were co-transfected with Rev-NLUC and CLUC-Rev plasmids and either empty vector, M4, M7, or wt Rev, followed by cell lysis and luciferase assay at 72 h. d) Similar to 4c) but with Rev mutants V16D, L60R, I55H, R48S, and wild-type. Student’s t-tests were performed to compare the RLU values resulting from the addition of each of the mutants relative to that produced by transfection of empty vector; symbol key is shown in Figure 2b. e) Each NLUC and CLUC plasmid as indicated was transiently transfected pairwise into 293T cells, and RLU measured after cell lysis at 72 h. Here, Ncontrol is N-FRB and Ccontrol is C-FKBP.

Host factor Crm1 binds to Rev that is multimerized on the RRE in order facilitate nuclear export of intron-containing HIV RNA. Previously we had constructed HA-tagged versions of both human (h) and mouse (m) Crm1; for these experiments we also made HA-tagged constructs of hCrm1 and mCrm1 that contained only the first ~500 amino acids by PCR amplification.

After confirmation of constructs by limited sequencing, these were transiently transfected into 293T cells to verify expression by immunoblotting using anti-HA antibody. Full-length hCrm1 and mCrm1 are known to be ~100 kDa (see Figure 3 of [51]), whereas the truncated forms were both ~50 kDa (Figure 3a). These plasmids were each transiently transfected into 293T cells in increasing amounts along with Rev-NLUC and CLUC-Rev and RLU measured at 72 h after cell lysis. We observed no significant reduction of RLU for any of the four Crm1 constructs, suggesting that Crm1 is unable to compete or modulate the Rev-Rev interaction in this system (Figure 3b).

**Figure 3.**
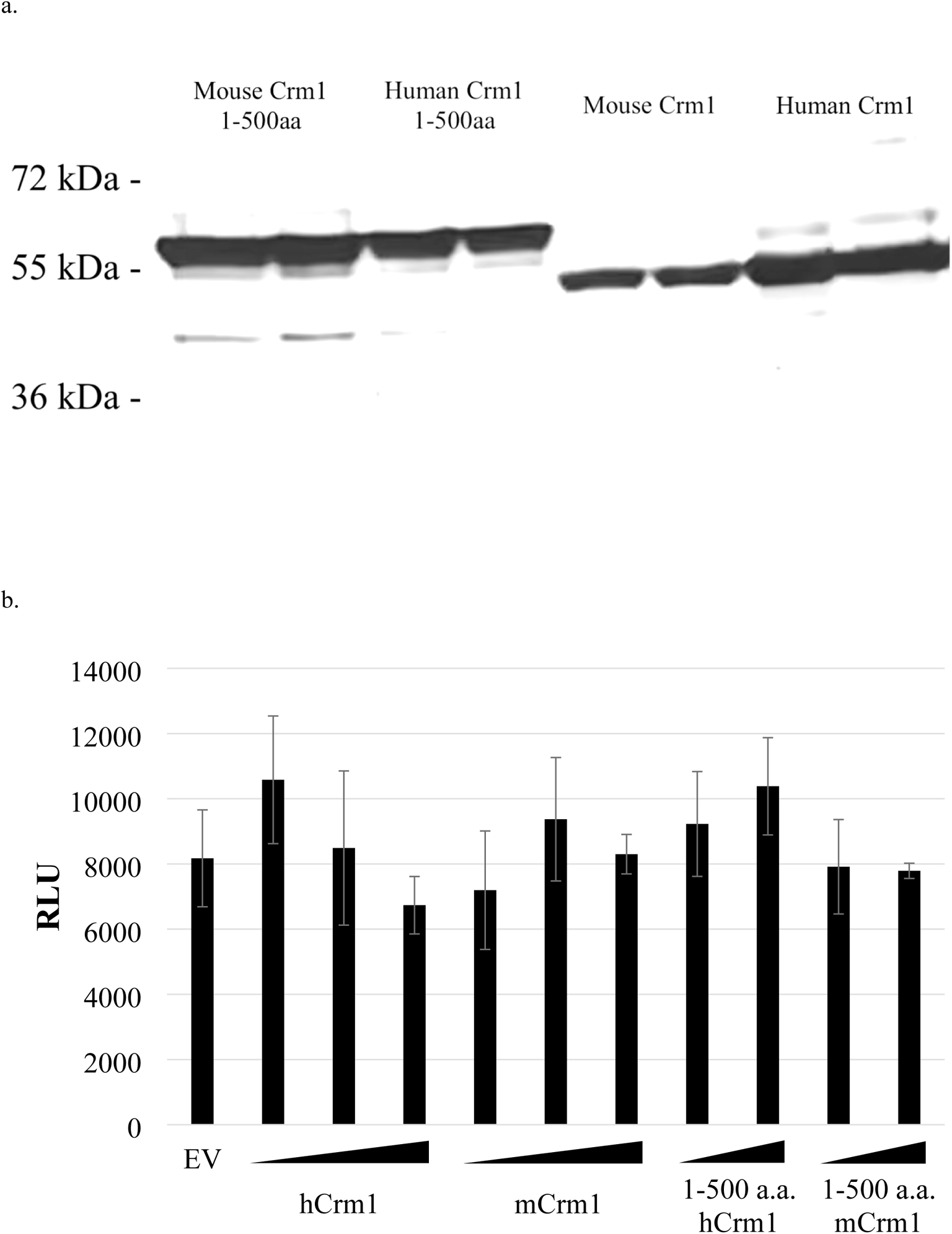
Additional Crm1 does not modulate RLU. a) Transfection of 293Ts was performed in duplicate with plasmid indicated at top of each lane. Immunoblot using anti-HA antiserum. b) 293Ts were transfected with Rev-NLUC and CLUC-Rev and either 4 μg of empty vector (EV, first bar), 1-2-4 μg of hCrm1 (first wedge), 1-2-4 μg of mCrm1 (second), 1-2-4 μg of 1-500 aa hCrm1 (third), or 1-2-4μg of 1-500 aa mCrm1 (fourth), lysed at 72 h, and assayed for luciferase activity. All plasmid constructs were CMV-driven. Values are mean ±SD. Student’s t-tests were performed comparing each experimental mean against empty vector, and in no case was P<0.05, with correction for multiple comparison testing.

### Testing Rev mutants in the SLCA

A number of Rev mutants have been characterized over the years, many of which are loss-of-function. Several have been identified as being unable to form multimers on radiolabeled RRE. We obtained M4 (SIS→I, aa 54-56) and M7 (YSN→DDL, aa 23,25,26) Rev, along with V16D, R48S, I55H, and L60R. Of note, in the original publication I55N [52] was reported but the plasmid sent to us was I55H. Although all of these mutant Revs had been previously characterized biochemically and functionally, we decided to test them in a quantitative assay for virus production. To do so, we established a Rev-complementation assay by first filling in 4 bases at a BamH1 site within the second exon of Rev in our HIV packaging vector or HIV-PV, thus frame-shifting and inactivating Rev. We co-transfected into 293T cells this HIV-PVΔRev, VSV G, a third generation HIV vector encoding *bsd^r^* gene, and each, unfused Rev plasmid.

Regarding the latter, although we had received M4 and M7 Revs in a CMV expression plasmid, for the other four mutant Revs we had to subclone each into CMV expression plasmid pCI (Promega). At 72 h, we collected transfected cells, size-separated RIPA lysates by SDS-PAGE, and performed immunoblotting using anti-Rev antisera. Wt and each of the mutant Revs were detected at ~18-kDa, although there was slight variability in protein size (Figure 4a). Of note, we could not reliably detect M7 Rev using available antisera, although we used an original plasmid construct from Bryan Cullen (Duke) which was sequenced to be correct. Cell culture supernatant was also harvested at 72 h and titered on 293T targets by serial dilution and enumerating blasticidin-resistant CFU. There was a reasonably good correlation between viral titer and RLU for each of the Revs tested (R=0.81; Figure 4b).

Full-length versions of all of these mutant Revs were PCR-amplified, sequenced confirmed, and each was separately cloned into the parental FRB-NLUC and CLUC-FKBP plasmids, as described above. We confirmed expression of each, again by transiently transfecting the plasmids into 293T cells followed by RIPA lysis of the cells at 72 h and SDS-PAGE/immunoblotting using anti-FFLUC antisera. The sizes of all the Rev mutant NLUC and CLUC fusions were approximately 70 kDa and 35 kDa, respectively, as expected (Figure 1a-d).

These Rev mutants were then paired and transiently co-transfected into 293T cells, with cell lysis and RLU measured at 72 h, normalized to wt Rev. For this experiment we had also made ‘empty vector’ or EV versions of FRB-NLUC and CLUC-FKBP by fully deleting FRB and FKBP, respectively, since we had occasionally observed some resultant RLU even in the absence of rapamycin for the parental plasmids. In rank order of RLU activity, lowest to highest, were EV, L60R, M7, R48S, I55H, V16D, and M4 (see x-axis of Figure 4b). Three had <25% of wt activity, whereas three had between ~25% and 35% of wt Rev activity. This suggests that all of the tested Rev mutants had impairment in terms of homologous interaction in this assay.

We next tested whether any of the mutant Revs could inhibit wt Rev interaction in the SLCA. Each Rev mutant in the CMV expression plasmid was transiently transfected into 293Ts, along with Rev-NLUC and CLUC-Rev, and RLU measured at 72 h after cell lysis. Only transfection of wt Rev significantly reduced RLU by 50% or greater (Figure 4c-d). We were also interested in determining whether mutant Rev-NLUCs and CLUC-Revs, when paired with wt Rev-NLUC and CLUC-Rev, could restore RLU. For this we focused on Rev mutants M4 and M7 and also used parental FRB-NLUC and CLUC-FKBP plasmids as additional controls. Each NLUC and CLUC plasmid was transiently transfected pairwise into 293T cells, and RLU measured after cell lysis at 72 h. Not surprisingly, parental FRB-NLUC+CLUC-FKBP and wt Rev-NLUC+CLUC-Rev gave the highest RLU, whereas all combinations involving M4 and M7 NLUC and CLUC plasmids resulted in much lower RLU (Figure 4e). Wt Rev-NLUC or CLUC-Rev could not rescue any of the M4 or M7 constructs. This is entirely consistent with the known loss-of-function of both M4 and M7 Revs and their failure to multimerize on the RRE.

### Anti-Rev nanobody increases RLU in the SLCA

Several years ago, a llama-derived, single chain only antibody that blocks HIV replication by targeting the N-terminal multimerization domain of Rev was isolated [53]. This nanobody (NB) inhibited Rev function but at the same time enhanced Rev dimerization. We obtained plasmid constructs of a control NB or the anti-Rev NB, each fused to Kusabira Orange (KO). The nature of the NB-KO fusions keeps the proteins from being secreted such that they remain in the cytoplasm (see Figure 5b). We confirmed that co-transfection of increasing amounts of the anti-Rev NB-KO along with HIV-PV, VSV G, and HIV-TV FG12 significantly inhibited vector production by ~ 94% compared to control NB-KO, whereas co-transfection of increasing amounts of dominant negative RevM10-eGFP significantly reduced vector production by >99% (Figure 5a). Next, wt Rev-NLUC and CLUC-Rev, along with increasing amounts of Rev-GFP, were transiently transfected into 293T cells with each NB-KO plasmid, and at 72 h cells were lysed and RLU measured. An aliquot of cells was subjected to flow cytometry to confirm ~50% transfection of the KO fusion, as assessed by PE-TX Red fluorescence (Figure 5b, right panels). The anti-Rev NB significantly increased RLU roughly 4-6 fold compared to control NB, and in both cases RLU were inhibited by increasing amounts of Rev-eGFP (Figure 5c-d). The observed increase in RLU was dependent upon transfection of both Rev-NLUC + CLUC-Rev (Figure 5c). Thus, the anti-Rev NB inhibited virus production and yet enhanced Rev-Rev interaction, as assessed by the SLCA, consistent with published results.

**Figure 5.**
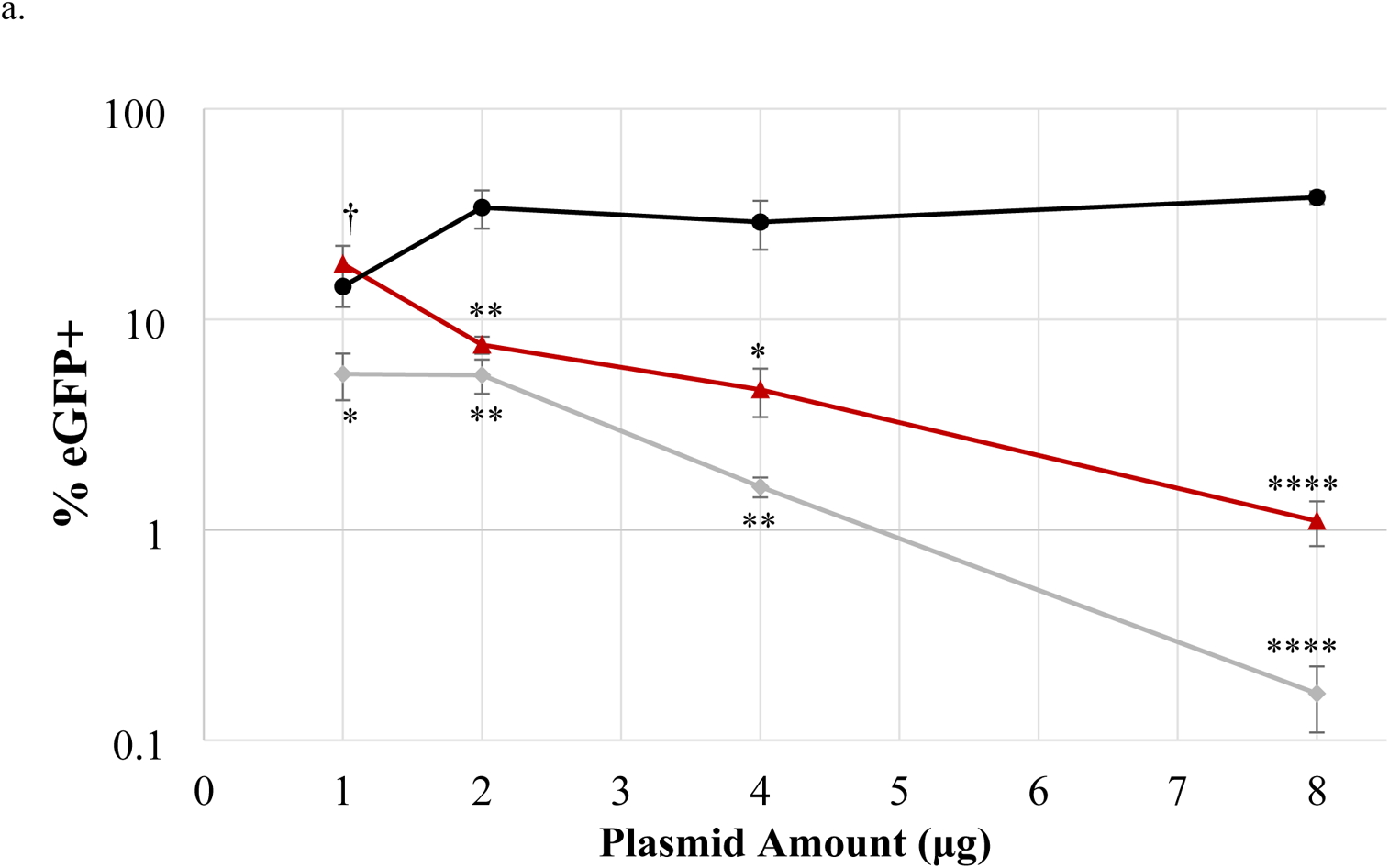
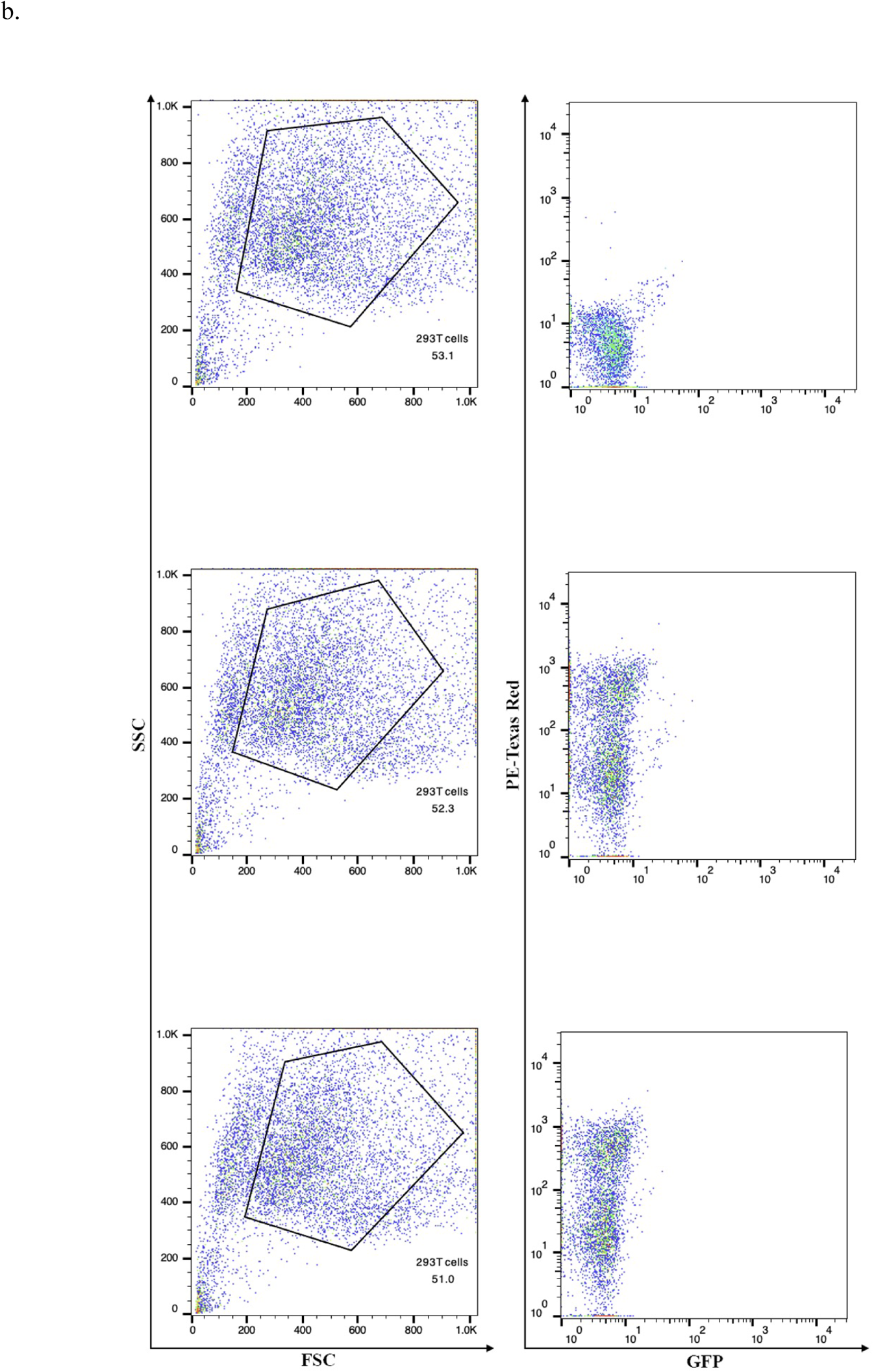
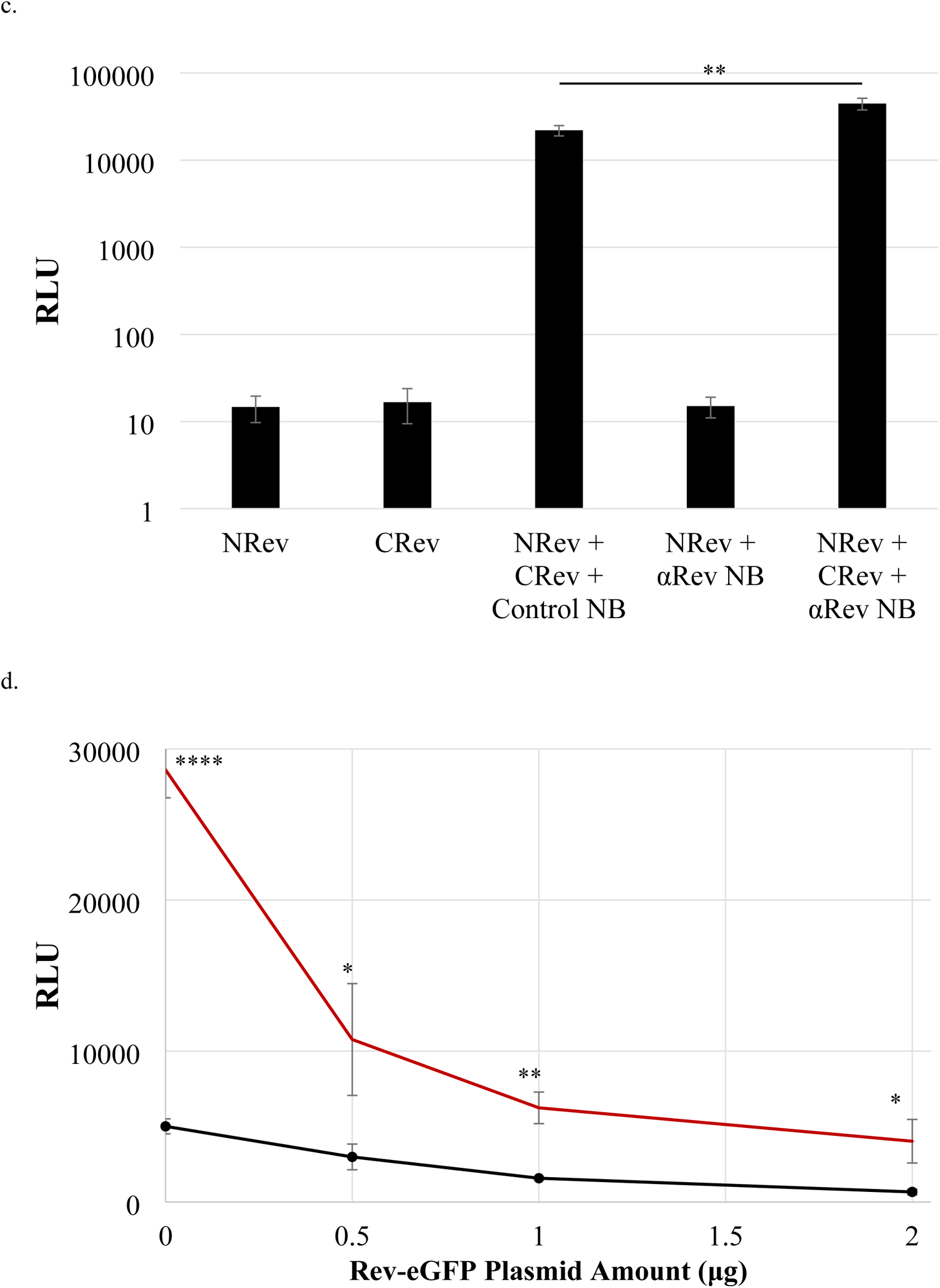
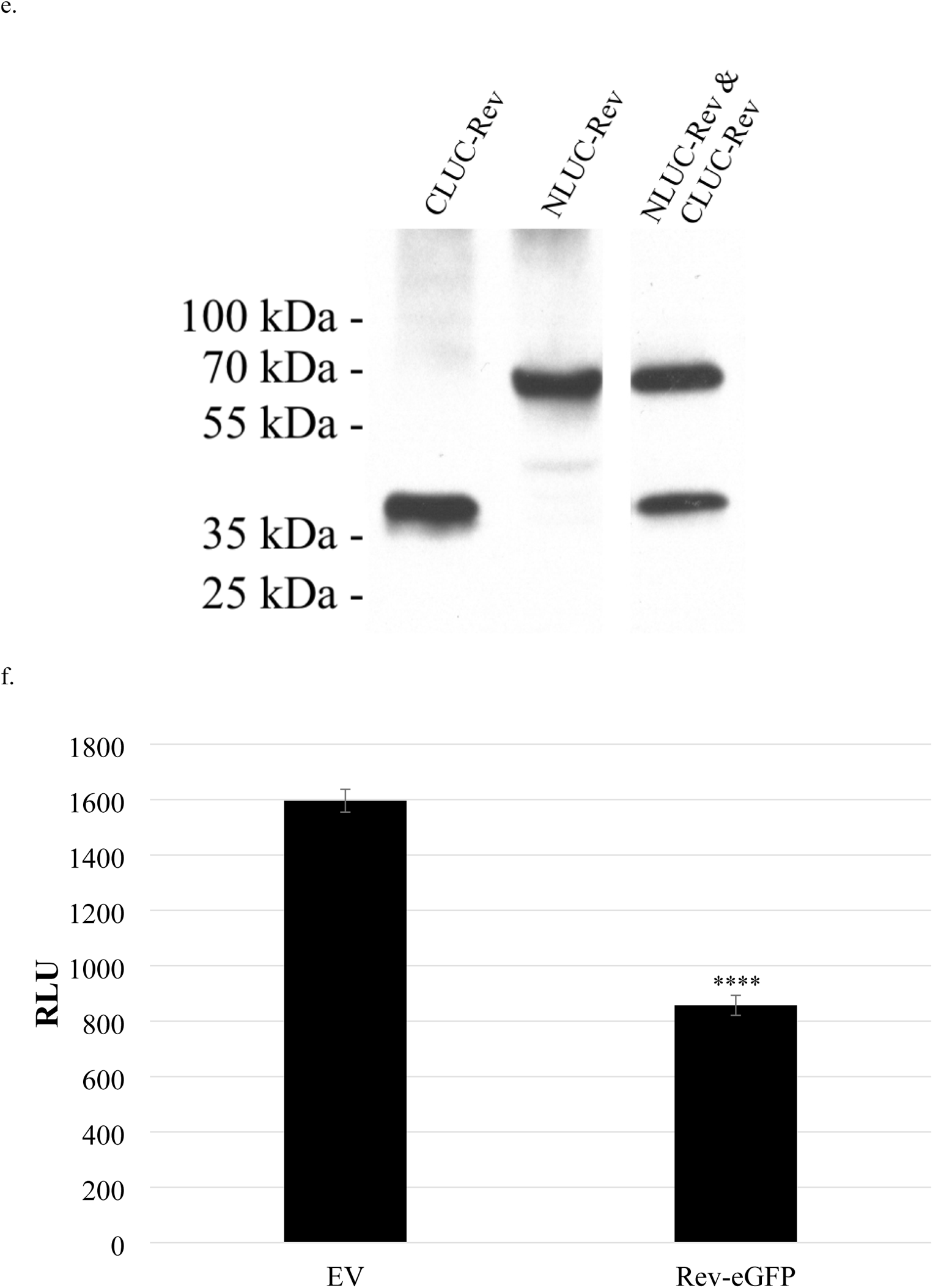
Effects of anti-Rev NB on virus production and RLU. a) 293Ts were co-transfected with increasing amounts of anti-Rev NB-KO (red triangles), control NB-KO (black circles), or RevM10-eGFP (grey diamonds) along with HIV-PV, VSV G, and HIV-TV FG12. At 72 h culture supernatant was harvested and titered on 293Ts, with readout being %eGFP+ cells, as measured by flow cytometry after another 72 h. Student’s t-tests were performed to compare the %eGFP+ values resulting from the anti-Rev NB and RevM10-eGFP against the control NB at each plasmid amount. b) 293T cells were either mock-transfected, or transfected with anti-Rev NB-KO or control NB-KO and subjected to flow cytometry at 72 h. Left panels are forward-side scatter with gating on viable cells indicated; right panels are PE-TX red vs. GFP. Top: mock transfection, middle: control NB-KO fusion, bottom: anti-Rev NB-KO fusion. c) Rev-NLUC and CLUC-Rev were transiently transfected into 293T cells along with indicated NB-KO plasmid, and at 72 h cells were lysed and RLU measured. d) Rev-NLUC and CLUC-Rev were transiently transfected into 293T cells with along with anti-Rev NB-KO (red squares) or control NB-KO (black circles) and increasing amounts of Rev-eGFP plasmid, and at 72 h cells were lysed and RLU measured. e) Lysates from stable 293T-based cell lines, expressing either CLUC-Rev, Rev-NLUC, or both, were immunoblotted using anti-Luciferase antiserum. f) 293T cells from the stable 293T line expressing both CLUC-Rev and Rev-NLUC were plated in 96-well format and transiently transfected with either empty vector or Rev-eGFP. Transfection efficiency, as judged by inverted fluorescence microscopy scoring for eGFP+ cells was approximately 50%. At 72 h, cells were lysed and RLU quantified. Z’ = 0.69. Refer to Figure 2 for symbol key for multiple parts of this Figure.

### Stable Rev-NLUC + CLUC-Rev cell lines recapitulate transient transfection results

Based upon our results with the transient transfections, we decided to produce stable, 293T-based cell lines expressing the FFLUC complementation components. We separately introduced the Rev-NLUC gene linked to IRES-puro^r^ and the CLUC-Rev gene linked to IRES-hygro^r^ into third generation HIV-based vectors. These were converted into VSV G-pseudotyped particles and separately and sequentially introduced into 293T cells such that the cells expressing Rev-NLUC were puro^r^, those expressing CLUC-Rev were hygro^r^, and those expressing both were doubly resistant. After passaging, expression of both FFLUC components was confirmed by RIPA lysis, SDS-PAGE, and immunoblotting using anti-FFLUC antisera (Figure 5e). Non-ionic detergent lysates were made from each and RLU quantified, and only the cell line that was doubly resistant gave rise to RLU. In order to determine the suitability of using such a cell line to screen for inhibitors of Rev-Rev interaction, cells were plated in 96-well format and transfected with either empty vector or Rev-eGFP. Transfection efficiency, as judged by inverted fluorescence microscopy scoring for eGFP+ cells, was approximately 50%. At 72 h, cells were lysed and RLU quantified. There was roughly a 50% decrease in RLU in the presence of Rev-eGFP, and the calculated Z’ was 0.69 (Figure 5f). Although that Z’ may be acceptable in some instances because the standard deviations were very low, the 2-fold dynamic range is not ideal for a high throughput screen.

### Mixing Rev-NLUC and CLUC-Rev cell lysates reproduce the cell-based results

Since working with intact cells for a high throughput screen would be inconvenient and logistically difficult, we decided to test whether mixing separate Rev-NLUC and CLUC-Rev cell lysates would give rise to RLU. For this, we used the stable cell lines described above, made non-ionic detergent lysates of each, mixed them together in equal amounts, and quantified RLU 1 h later. Individually, Rev-NLUC and CLUC-Rev lysates produced relatively low RLU, whereas upon lysate mixing resultant RLU increased by 100-1000 fold (Figure 6a). In addition, we quantified RLU at different times after lysate mixing. Within minutes, RLU were produced, and values were relatively stable for 60 minutes after mixing, with a slight significant increase in RLU at 60 min (Figure 6b). Furthermore, we transiently transfected the Rev-NLUC stable cell line with EV or Rev-eGFP, made lysates at 72 h, and mixed each with CLUC-Rev lysate. Even after 1 minute RLU was reduced when Rev-eGFP had been transfected, with marginally greater inhibition after 60 minutes (Figure 6c). These results suggest that the cell-free Rev-Rev interaction occurs very rapidly, and inhibition by excess Rev is also quite fast, although the latter may be due to pre-formed Rev multimers in the cells.

**Figure 6.**
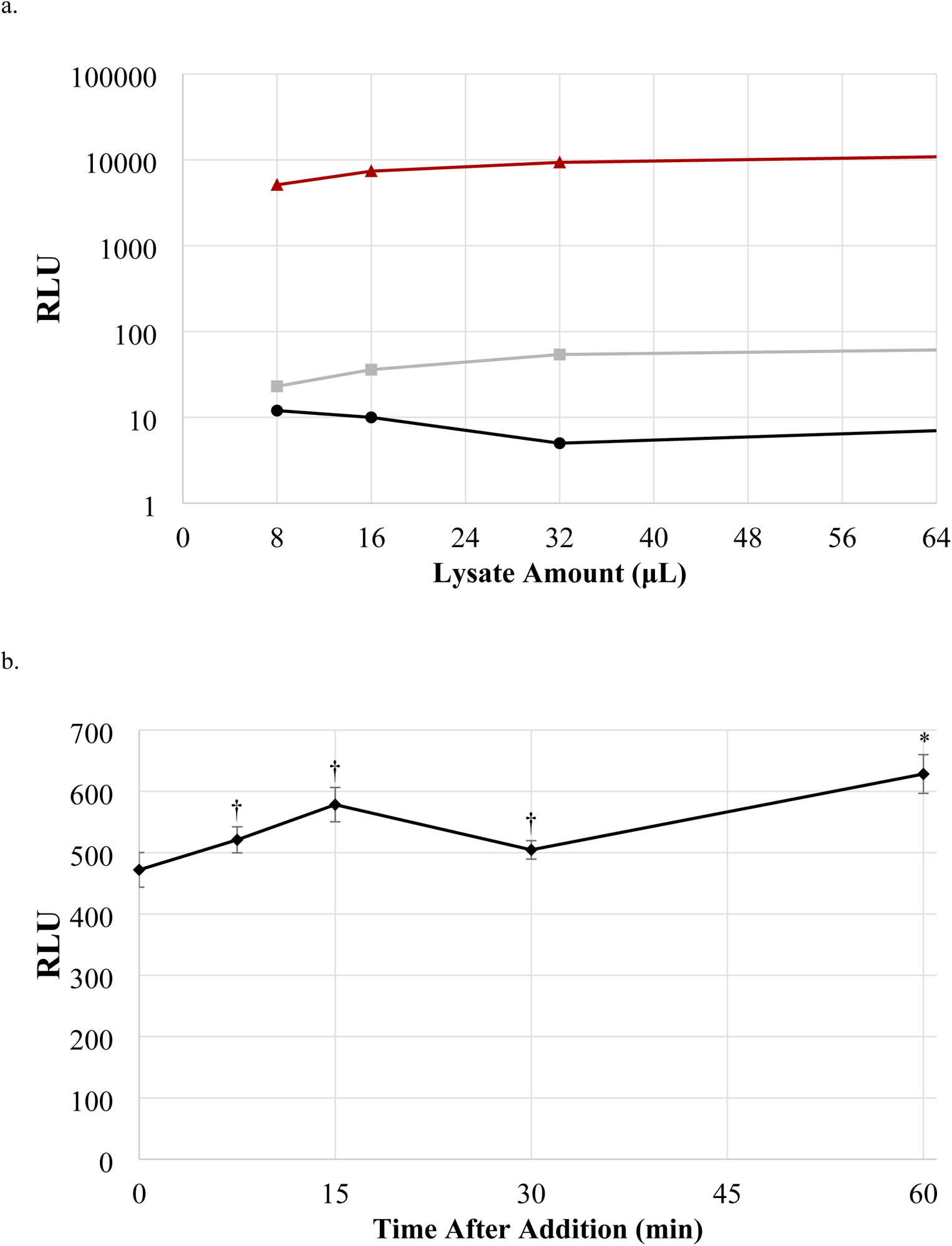
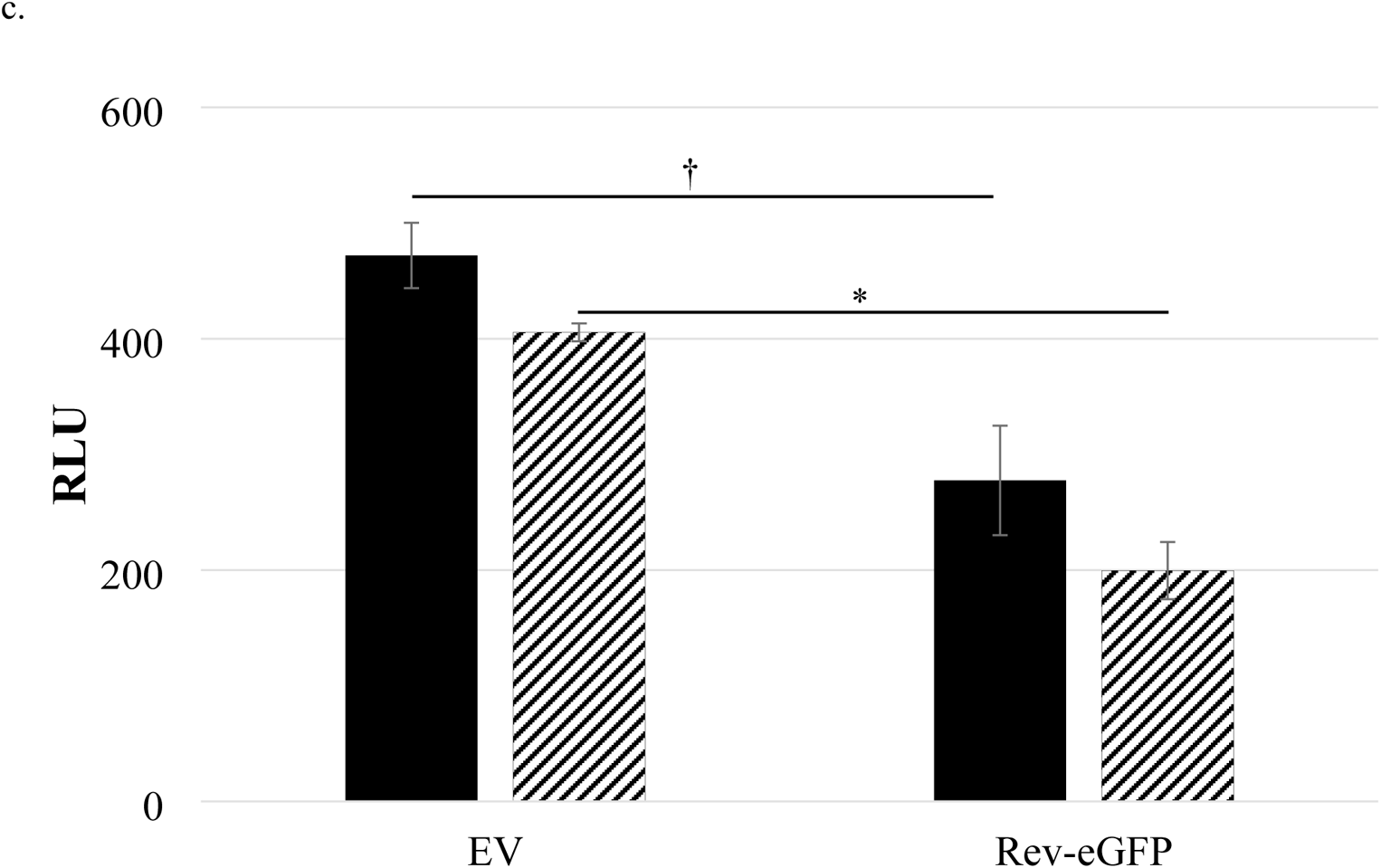
Mixing separate Rev-NLUC and CLUC-Rev cell lysates yields RLU. Non-ionic detergent cell lysates were prepared from the stable 293T cell lines separately expressing Rev-NLUC (black circles) and CLUC-Rev (grey squares). Lysates were mixed in equal amounts, and RLU quantified 1 h later (red triangles). b) Lysates from part a) were mixed and RLU measured at increasing times. RLU values remained high and relatively stable. c) The Rev-NLUC stable cell line was transiently transfected with empty vector (EV) or Rev-eGFP, lysates made at 72 h, and each mixed with CLUC-Rev lysate. RLU measured after 1 (black bars) or 60 minutes (striped bars). Refer to Figure 2 for symbol key for multiple parts of this figure.

### Further characterization of the cell-free Rev-Rev interaction

We wished to test whether any of the Rev mutants had activity in the cell-free assay. NLUC and CLUC Rev mutants were individually transiently transfected into 293T cells and lysate produced at 72 h and then mixed at 1:1 ratio. As anticipated, wt Rev-NLUC mixed with wt CLUC-Rev had the greatest FFLUC activity, whereas the Rev mutants had varying RLU levels that were lower, fairly consistent with the cell-based assay (Figure 7). Of note, no mutant combination had RLU as high as wt-wt, suggesting that the mutant Revs could not trans-complement each other in this assay, with the possible exceptions of CLUC-Rev L60R and M4 Rev-NLUC, CLUC-Rev V16D and V16D Rev-NLUC, and CLUC-Rev and V16D Rev-NLUC pairs.

**Figure 7.**
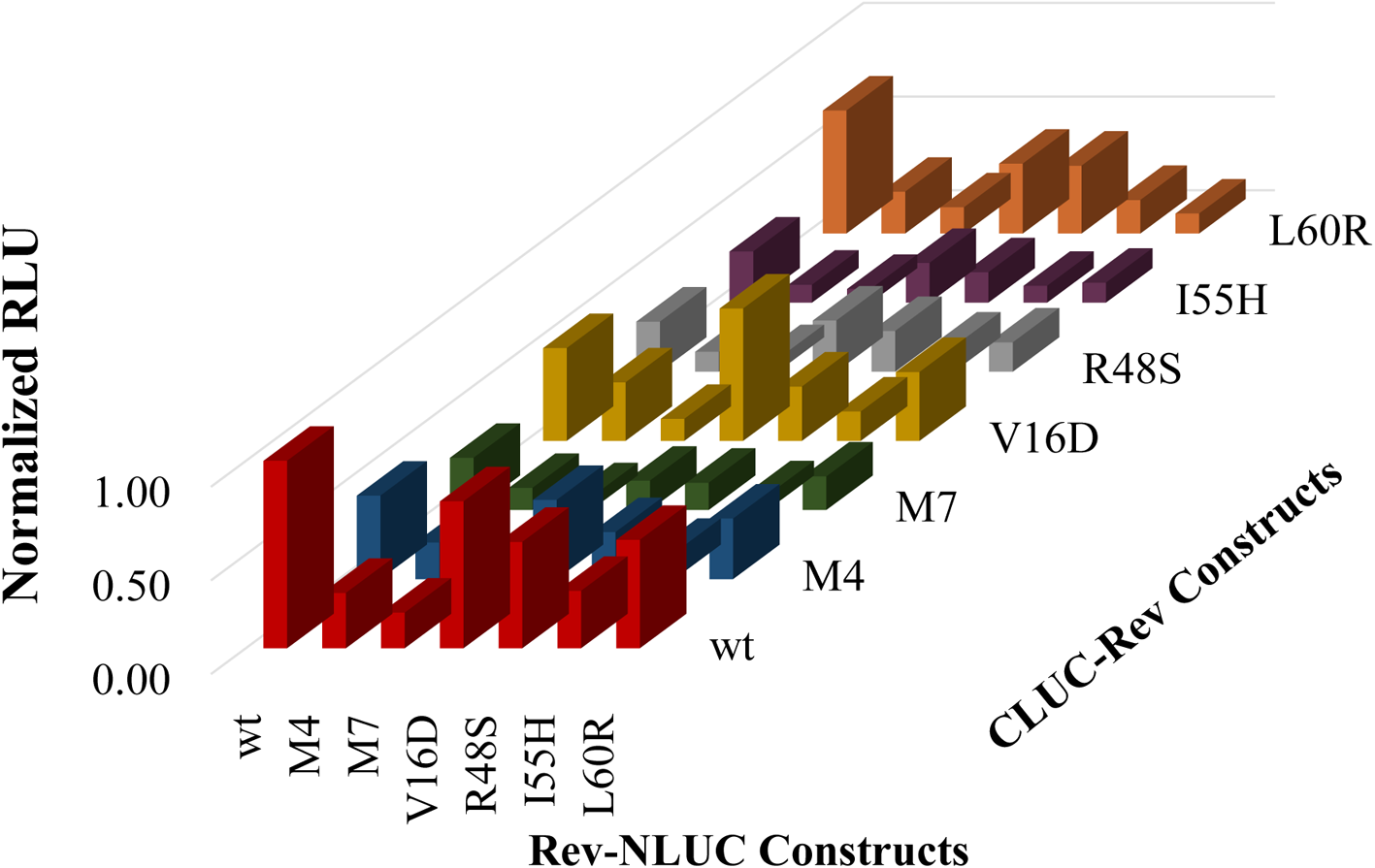
RLU activity of wt and mutant Rev-NLUC and CLUC-Rev pairs. Indicated wt or mutant Rev-NLUC and CLUC-Rev plasmids were individually transfected into 293T cells, with cell lysis at 72 h. RLU was then quantified of the 49 combinations of Rev-NLUC and CLUC-Rev lysate pairs after 1 h incubation and normalized to the wt/wt pair (set at 1.0).

Treatment of lysates made from the stable wt Rev-NLUC and CLUC-Rev cell lines with DNAse for 30 min prior to mixing had no effect on subsequent measurements of RLU activity, whereas similar treatment with RNAse slightly but significantly reduced RLU at the two lower lysate amounts (Figure 8a). Because these stable cell lines were transduced with an HIV vector that has an intact RRE, we thought it would be of interest to investigate whether addition of RRE RNA by itself would modulate activity in the cell-free lysates. Control or RRE RNA synthesized in vitro was added to lysates made from the stable wt Rev-NLUC and CLUC-Rev cell lines and RLU measured 30 minutes later. In the presence of HIV RRE RNA specifically RLU nearly doubled, which was significant (Figure 8b), consistent with the RRE having a positive influence on Rev-Rev interaction. We wished to test whether the anti-Rev NB would also enhance RLU in the cell-free assay. The control or anti-Rev NB-KO fusion gene was cloned into a third generation HIV vector downstream of the CMV promoter and upstream of IRES-*bsd^r^* cassette and each separately introduced into stable, puro^r^ wt Rev-NLUC 293Ts by VSV G-mediated transduction, resulting in a pool of bsd^r^ + puro^r^ cells that were also KO+, as judged by flow cytometry. Lysate prepared from wt CLUC-Rev stable 293T cells was mixed with that of Rev-NLUC+control NB-KO or Rev-NLUC+anti-Rev NB-KO and RLU measured 30 min later.

**Figure 8.**
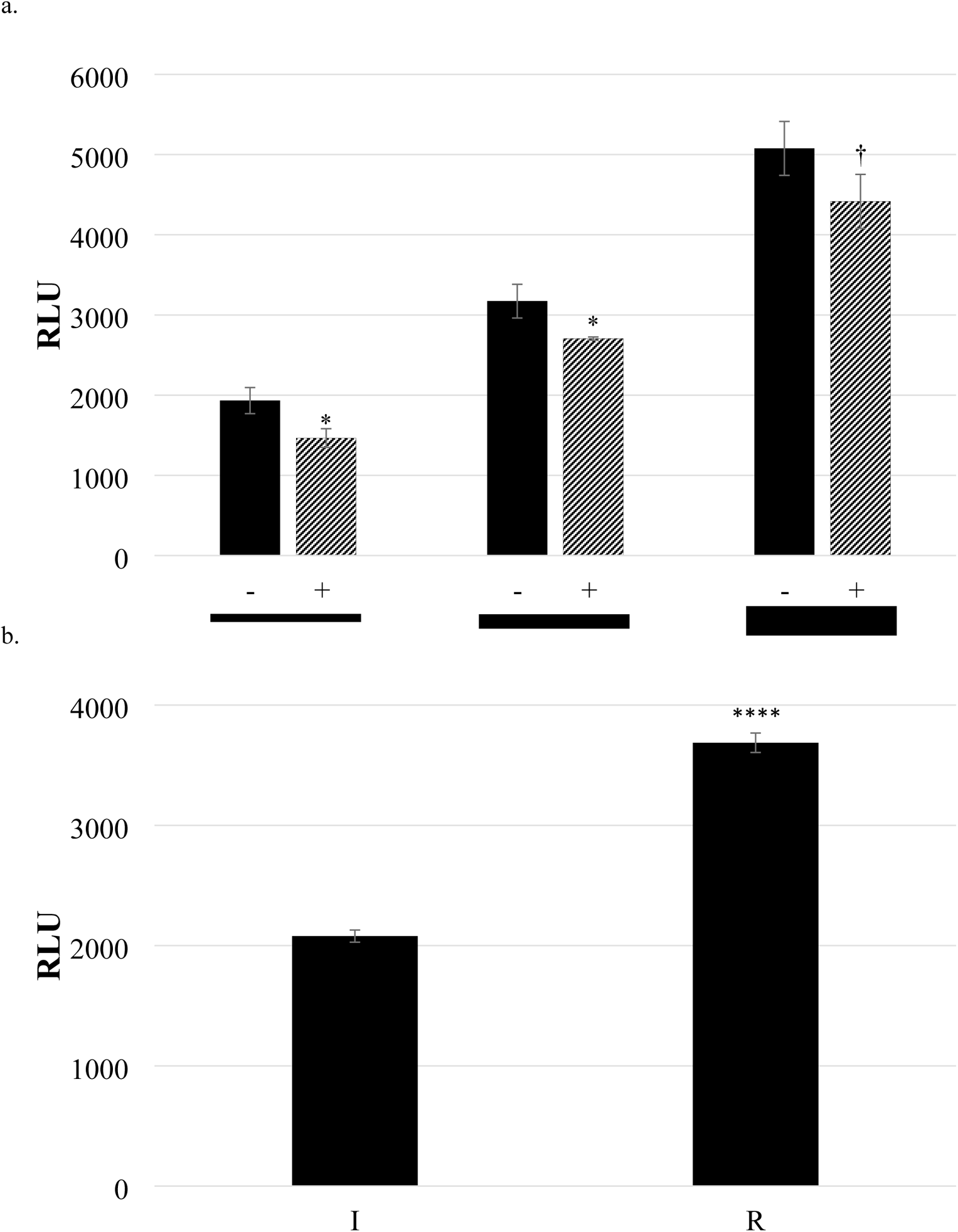
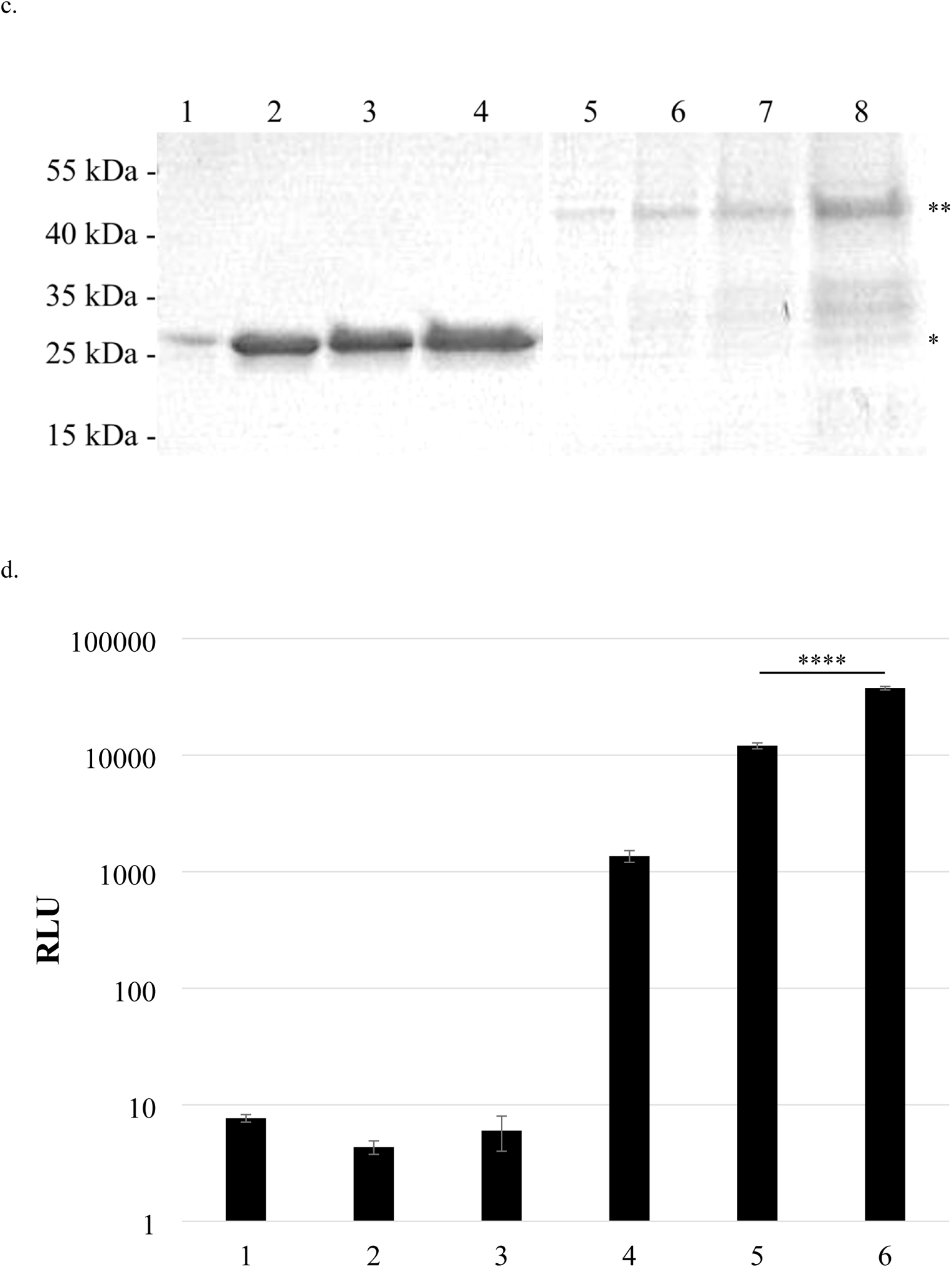
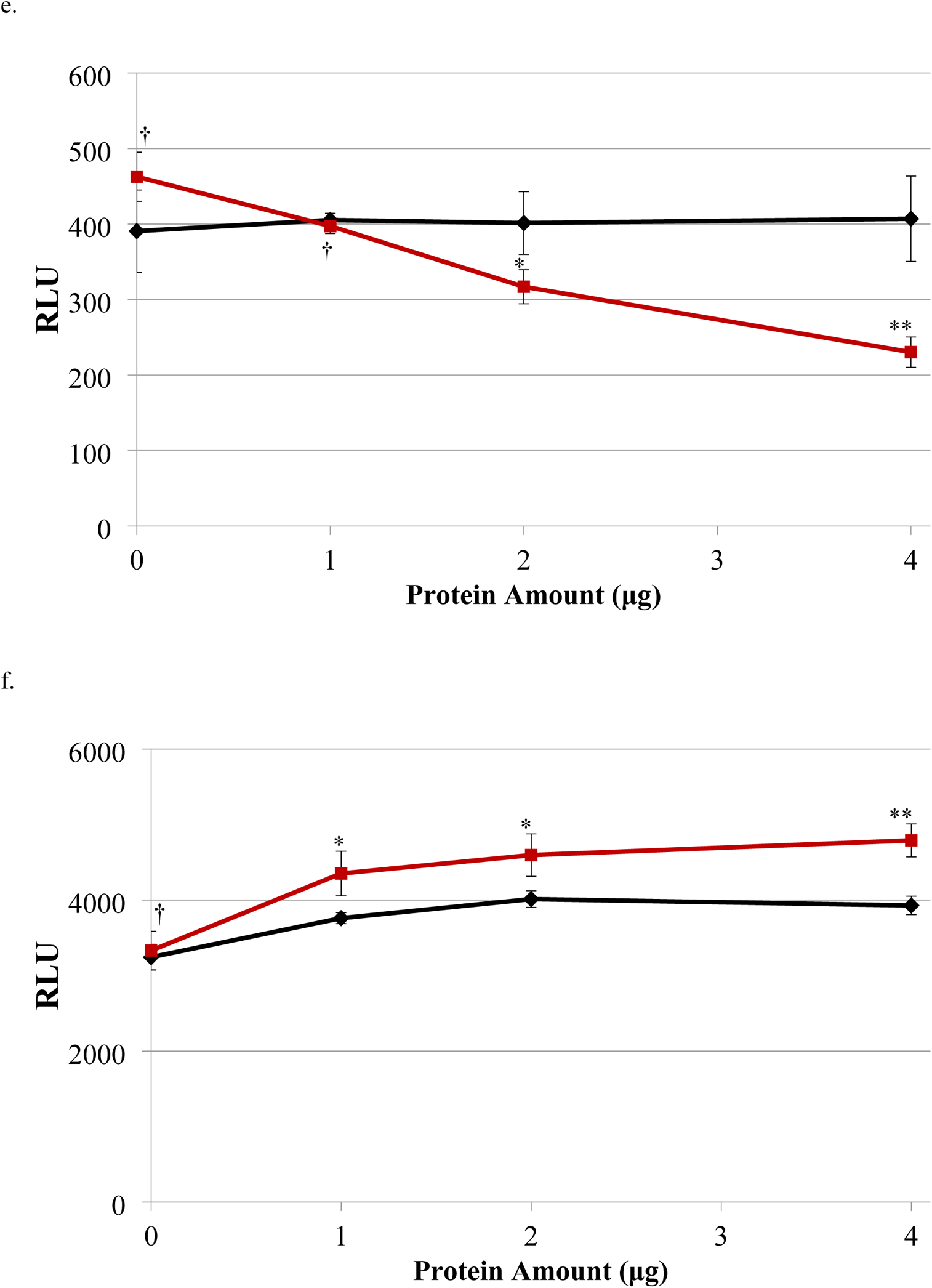
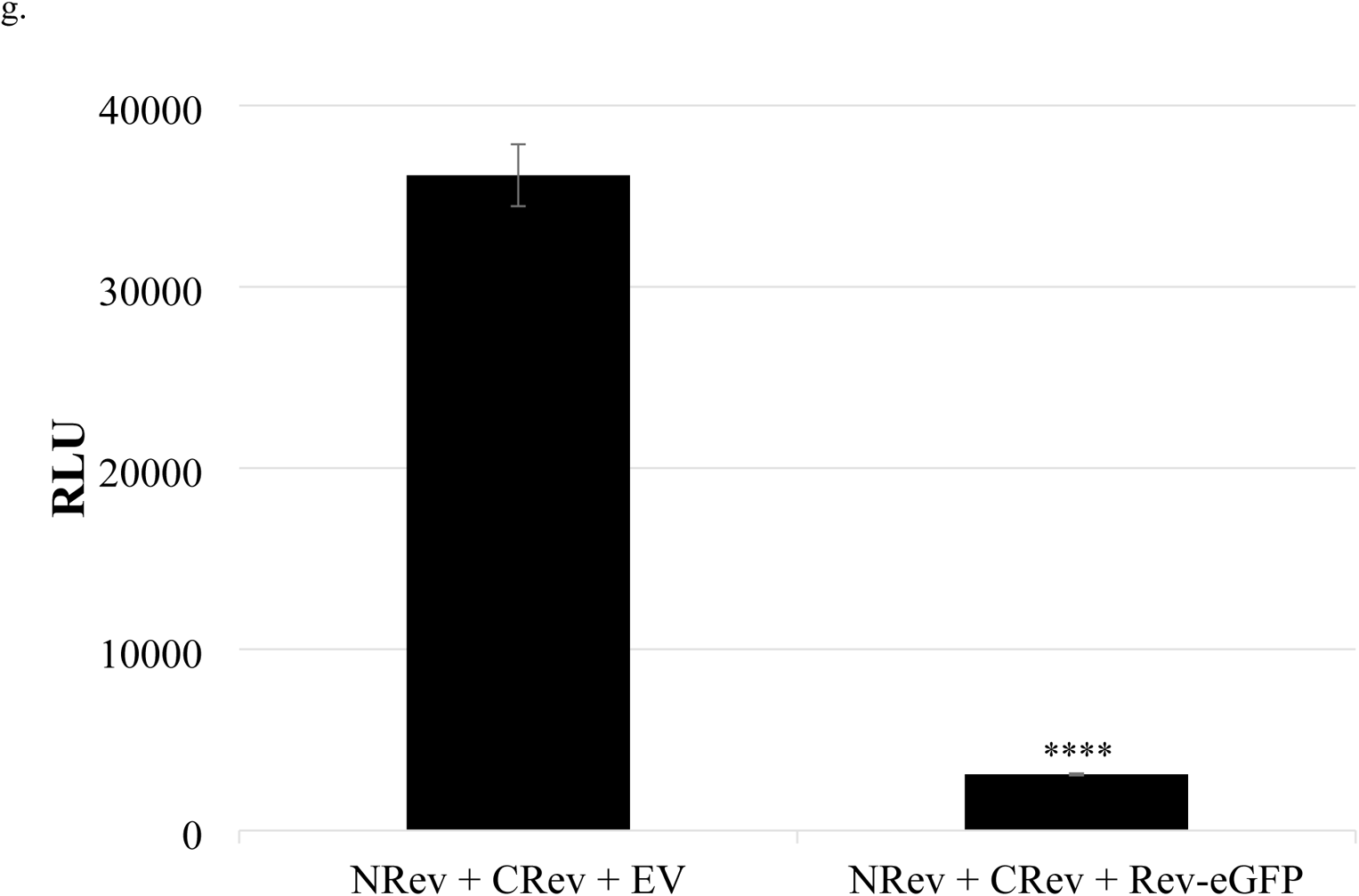
Characteristics of the cell-free Rev-NLUC and CLUC-Rev interaction. a) Lysates from Rev-NLUC and CLUC-Rev stably expressing 293T cells were separately prepared, mixed, and treated with 10μg of RNAse A (+) or buffer (-) in triplicate for 30 minutes prior to luciferase assay. Shown are mean values ± SD. Thickness of red bar below indicates relative amount of lysate used. Student’s t-tests were performed to compare the RLU values that resulted in the presence and absence of RNAse A all three lysate amounts. b) Lysate from stable Rev-NLUC expressing cells was added to 10 ng irrelevant RNA (I) or 10 ng 325 base HIV RRE RNA (R), then lysate from stable CLUC-Rev expressing cells was added. Both RNAs had been in vitro transcribed using linearized plasmid template with SP6 phage RNA polymerase and intactness confirmed by gel electrophoresis. After 30 minutes RLU was measured. Shown are mean values ± SD. c) Coomassie Blue stained protein gel of increasing amounts of GST alone (lanes 1-4) or GST-Rev (lanes 5-8). d) Lysates prepared from 293Ts stably or transiently expressing indicated genes were mixed and RLU measured 30 min later. Bars: 1: mock transfection, 2: CLUC-FKBP (transient), 3: FRB-NLUC (transient), 4: CLUC-FKBP + FRB-NLUC (transient), 5: Rev-NLUC/control NB-KO + CLUC-Rev (stable), 6: Rev-NLUC/anti-Rev NB-KO + CLUC-Rev (stable). Note log scale for RLU. e) and f) Cell-free Rev-Rev interaction inhibited by GST-Rev. GST-Rev fusion protein (red squares) and GST alone (black diamonds) were purified from E. coli. Each was added in increasing amounts to lysates of 293Ts separately stably expressing Rev-NLUC and CLUC-Rev (d) or CMV-FFLUC (e) at the time of mixing and RLU measured 30 min later. Z’ was <0.5. g) Cell lysates from 293Ts stably expressing either Rev-NLUC, Rev-NLUC/Rev-eGFP, or CLUC-Rev were separately prepared and mixed as indicated in 96-well format, with RLU measured 30 minutes later. Values reflect those of 10^5^ cell equivalents. Calculated Z’ ~0.85. Statistical testing was performed using student’s t-test with correction for multiple comparison testing in some cases; see Figure 2 for symbol key for multiple parts of this figure.

Similar to the cell-based system, there was a significant, 3-fold increase in RLU in the presence of the anti-Rev NB-KO gene, suggesting that it can enhance Rev-Rev interaction even in a cell-free system, after simply mixing lysates (Figure 8c).

### Cell-free Rev-Rev interaction is inhibited by excess Rev

In order to determine the Z’ for the cell-free system we first purified 45 kDa GST-Rev fusion protein and GST alone from E. coli (Figure 8d). The GST-Rev was contaminated with some breakdown products. This was added in increasing amounts to wt Rev-NLUC and CLUC-Rev lysates at the time of mixing and RLU measured 30 min later. GST-Rev, but not GST alone, significantlyinhibited RLU production at the higher amounts used (Figure 8e). Reduction of RLU was not observed for either GST-Rev or GST alone after addition to lysates made from 293T cells that had been transfected with a CMV-FFLUC reporter (Figure 8f). If anything, addition of GST-Rev resulted in marginally higher RLU. Although these results suggest that the observed decrease in RLU is specific to the Rev-Rev interaction, the degree of inhibition observed was only two-fold, and the resultant Z’ was <0.5.

This suboptimal Z’ may be due to the fact that most of the GST-Rev purified from E. coli was not folded properly and thus would not inhibit Rev-Rev interaction. In order to generate a system in which most of the Rev is functional, we decided to over-express Rev-eGFP in Rev-NLUC-expressing 293T cells. To do so, Rev-eGFP was inserted downstream of CMV promoter and upstream of an IRES-bleo^r^ cassette in a third generation HIV vector. This vector was introduced by VSV G-mediated transduction into Rev-NLUC puro^r^ 293T cells and selected in both phleomycin and puromycin. In the absence of any cell sorting, the majority of the resultant stable 293T cells were eGFP+, based upon flow cytometry. Lysates from these 293T cells were prepared and mixed with those from the CLUC-Rev stable 293T cells and RLU measured 30 min later. In comparison to simply mixing CLUC-Rev and Rev-NLUC 293T cell lysates, presence of Rev-eGFP significantly reduced RLU >90%, with resultant Z’ of ~0.85 (Figure 8f). This degree of RLU inhibition suggests that Rev-eGFP multimerized with Rev-NLUC in the intact cells, prior to cell lysis.

## Discussion

Rev is an essential, highly conserved regulatory HIV gene, required for export of unspliced and partially spliced HIV transcripts from the cell nucleus to the cytosol, in order to produce Gag, Pol, and Env, but also to allow packaging of full-length, proviral RNA into viral particles that are assembled at the plasma membrane. In the absence of Rev the virus cannot replicate, and no artificial, man-made system has fully circumvented this gene requirement.

Although Rev does not have any known enzymatic activity and mediates its replicative function by binding to other Rev proteins, specific host factors, and the RRE present on these RNAs, it would be an ideal target for antiviral therapeutics. This is especially true because there are no cellular homologs of Rev or the RRE. Unfortunately, despite years of study, no inhibitor of Rev function has ever entered late-stage clinical trials and no anti-Rev compound, molecule, or gene has been approved by the FDA or any other regulatory agency for therapeutic use.

Possible reasons for this failure include the following: i) Rev does not have an easily assayable biochemical function that lends itself to a high throughput screen, ii) no high-resolution structure of Rev multimers bound to RRE and host factors exists to allow rational, computational design of inhibitors, iii) ease of virological escape from an anti-Rev compound, and iv) there is no need or desire to identify such an inhibitor. In terms of the latter, although it is true that available today there are more than two dozen antivirals FDA-approved against HIV, several of which are offered as single daily doses and are safe and efficacious, because of drug toxicities and tolerabilities coupled with the continued threat of virus resistance, there remains a persistent need for anti-HIV medications, especially those that target novel aspects of the replicative lifecycle, so called ‘first-in-class’ medications.

It should be also noted that for HIV+ individuals with suppressed viral loads on combination therapy, none of the medications block viral RNA transcription, RNA export from the nucleus, or release of viral particles from cells. This may explain why, using ultrasensitive assays, very low viral loads are detectable in these patients and in pre-clinical, nonhuman primate models and why cell activation using various latency reversal agents or LRAs can result in higher levels of plasma viral RNA [54–56]. Not only would an anti-Rev compound block viral replication overall, but it should further reduce the very low plasma circulating RNA levels in patients and also prevent increases in plasma RNA levels after treatment with LRAs and similarly acting compounds. Whether this would be comparable to a ‘block-and-lock’ approach is debatable, but use of an anti-Rev drug could shed light on the ongoing, controversial question of virus evolution in the setting of ART [57–63].

As a first step towards identifying such an anti-Rev therapeutic, herein we used SLCA to quantify Rev-Rev interaction, which is considered to be essential for Rev function. This system relies on the fact that the N and C termini of FFLUC, if separated, have virtually no enzymatic activity, but if brought close enough together by interacting proteins enzymatic activity is restored, at least to a certain degree. We demonstrated that this occurs when full-length Rev is fused to both the N and C termini of FFLUC and both are expressed in the same cell, whereas Rev mutants resulted in reduced FFLUC activity, highly correlated with other functional assays of the mutants. FFLUC activity was inhibited when wt Rev, and not mutant versions, were co-transfected, presumably through binding competition. The fact that the FFLUC activity was present in the soluble fraction of cell lysates strongly argues against it being a result of non-specific protein aggregation and instead that it is a true quantitative measurement of Rev-Rev interaction. Of note, this interaction occurred in the absence of RRE RNA.

Host factors Crm1 and Ran-GTP bind to Rev that is multimerized on the RRE to facilitate nuclear export of viral, intron-containing RNAs. Thus, it is conceivable that these host factors and perhaps others can enhance or stabilize the Rev-Rev interaction. Here, transfection of additional human or mouse Crm1 did not increase the interaction. In addition, expression of the first 500 amino acids of Crm1 from either species also did not modulate the interaction.

These results may be due to the fact that the cells already express abundant amounts of Crm1 or that Rev-Rev interaction is independent of Crm1. We did not try to reduce or knockout Crm1 or Ran gene expression since both are essential for cell viability and any results would be confounded by cytotoxicity. It should be noted that we do not know where in the cell these FFLUC-Rev fusion proteins are expressed, when the interaction occurs, the reversibility of the interaction, nor the true stoichiometry that results in FFLUC activity. Nor have we attempted to measure the K_D_ of the Rev-Rev association in this system, which could be confounder by multimerization effects.

Interestingly, the anti-Rev NB, which should be expressed in the cytosol, enhanced Rev-Rev interaction. Expression of this NB profoundly reduced virus production, as had been demonstrated previously, although in our hands it was not as effective as the trans-dominant Rev M10 mutant. Previous biochemical experiments had shown that the NB stabilizes Rev dimers and prevents formation of multimers on RRE RNA [53], but in the absence of a high-resolution structure its precise mechanism of action is uncertain. It is conceivable that it recognizes an epitope at the Rev-Rev interface, which then stabilizes the Rev dimer and yet impedes further multimerization, essential for Rev function. Another possibility is that NB binding to Rev prevents Rev from being shuttled in and out of the nucleus, inhibiting infectivity. That we observed greater FFLUC activity in the presence of the anti-Rev NB is further proof that we are truly quantifying a specific Rev-Rev interaction.

Although a stable cell line expressing both the NLUC and CLUC Rev fusions has a certain degree of utility, it would be far more advantageous to have a well-behaved cell-free system, especially for optimization of a high throughput assay in order to identify anti-Rev compounds. We were somewhat surprised that we were able obtain RLU simply by mixing lysates from cells that expressed each FFLUC component separately. The interaction occurred within minutes and was stable for over at least an hour. It was reduced slightly by treatment with RNAse but not DNAse, and addition of in vitro synthesized RRE RNA enhanced the interaction. We did not attempt to deplete or add additional purified Crm1 or any other host factor to the cell-free system. Even the different Rev mutants tested could interact in the cell-free system to varying degrees. We were only able to inhibit the cell-free interaction by adding purified GST-Rev or by co-expressing Rev-eGFP in one of the stable cell lines. It should be noted that the Rev-eGFP used in these experiments is the most functional Rev fusion we have identified thus far, by use of our Rev trans-complementation assay.

Would the cell-free system be amenable to a high throughput assay to quantify Rev-Rev interaction and subsequent disruption? First, we now have stable 293T-based cell lines that separately express the NLUC and CLUC components. These lines behave well and propagate easily, even in the presence or absence of selection, and their properties appear stable after more than one year of use and testing. Second, large volumes of cell lysate can be stored frozen indefinitely in the presence of 10% glycerol, and FFLUC activity is reconstituted after thawing and mixing. Third, the volume amount required to achieve >1000 RLU would be just a few μl of each cell lysate, making it amenable to 384 well screening format. One thousand RLU per 384 well would be approximately three orders-of-magnitude above background, and screening 100,000 compounds at a single concentration in triplicate would necessitate a total of 3 x 10^8^ RLU. We estimate this would require making lysate from ~one hundred 15 cm confluent plates of each cell line, which is easily achieved.

Perhaps a more pressing concern is the absence of a known small molecule inhibitor of Rev-Rev interaction. Although this is not an absolute requirement of a high throughput screen, it is difficult to reliably calculate a Z’ in its absence. In order to determine a Z’ here, we transduced one of the stable cell lines with Rev-eGFP, which markedly reduced resultant RLU after mixing the two cell lysates. The ten-fold dynamic range, coupled with the low standard deviations, presumably related to the fact that cell lysates were simply mixed in the absence of any other manipulation, gave a Z’ that in 96 well format was quite suitable for a high throughput screen. A larger, unanswered question is whether a small molecule would be capable of disrupting such a protein-protein interaction, especially when structural information is incomplete, as in the case here. There are now clear examples, however, of protein-protein interactions being hindered by small molecules, with functional consequences [64]. These compounds are now being advanced clinically, especially as anti-cancer therapeutics [65–68].

In summary, based upon the split-luciferase complementation system, we have developed a facile, rapid, and reliable assay that quantifies the ability of Rev to interact with itself. FFLUC activity is reconstituted in intact cells but also by mixing cell-free lysates. Although we were unable to modulate the interaction with cellular host factors known to interact with Rev, RRE RNA enhanced the cell-free association as did an anti-Rev NB. The cell-free system, which is based upon two stable cell lines, had the requisite characteristics desired of a high throughput assay capable of identifying inhibitors of Rev-Rev multimerization. Thus, employment of such an assay may allow for the discovery of first-in-class, anti-Rev compounds or small molecules that would complement the anti-HIV armamentarium.

## Methods

### Plasmids

Complementary (c) DNA of wt Rev from isolate NL4-3 was used as template to PCR amplify the 116 amino acid residue open reading frame or ORF using primers 5’-CCAGATCTATGGCAGGAAGAAGCGGAGACAGCG-3’ and 5’-GGTTACCGTACGCCTTCTTTAGTTCCTGACTCCAATACTGTAGG-3’. After cloning into pCR blunt II Topo vector (Invitrogen), the ~350 bp insert was sequenced, cleaved with BsiW1 and BglII, and the ~350 bp PCR product was cloned directionally into pFRB-NLUC (Addgene) cut with the BamH1 and BsiW. Because of an error in one of the primers, this plasmid was linearized with BsiW1, briefly treated with T4 DNA polymerase to remove four bp, and resealed to construct pRev-NLUC. pCLUC-Rev was similarly constructed, using primers 5’-GGCGTACGGGGCAGGAAGAAGCGGAGACACG-3’ and 5’-GGGATATCCTATTCTTTAGTTCCTGACTCCAATACTGTAG-3’ to amplify the Rev ORF, cleaved with BsiW1 and EcoRV, and inserted into pNLUC-FKBP (Addgene) precut with the same enzymes. The junction regions of both pRev-NLUC and pCLUC-Rev were sequenced to confirm intactness of the fusion ORF. cDNAs of Rev mutants M4 and M7, each in a CMV-driven expression plasmid, were obtained from Bryan Cullen of Duke University and those of Rev mutants V16D, R48S, I55H, and L60 were obtained from Joel Belasco of NYU. The ORFs of the latter four were separately blunt-cloned into the Sma1 site of pCI (Promega) to construct CMV-driven expression plasmids of each. To construct NLUC and CLUC versions of each Rev mutant, each was similarly PCR-amplified as wt, sequence-confirmed, and cloned. pHA-hCrm1 and pHA-mCrm1 were as described previously. To construct expression constructs of the first 500 amino acid of each, the forward primer used for each was 5’-GGCTCGAGGTCCCAGCAATTATGACAATGTTAGCAGACCATG-3’ and the reverse primers for hCrm1 and mCrm1 were 5’-GGGCGGCCGCTTATCCACTAATGGAGCCTATTGCCCAACAC-3’ and 5’-GGGCGGCCGCTTAGCCACTGATGGAGCCTATTGCCCAACAC-3’, respectively. These were used to PCR-amplify ~1500 bp product using the respective templates, which was first cloned into pCR blunt II Topo vector, sequenced-confirmed, then directionally cloned into pCMV-HA (Clontech) using the Xho1 and Not1 sites.

pGST-Rev was as previously described; pGEX-5X-2 encoding GST alone was purchased (Pharmacia). HIV RRE was PCR-amplified using primers 5’-GCAGAGAAAAAAGAGCAGTGGGA-3’ and 5’-CCAGGAGCTGTTGATCCTTTAGGTA-3’ and cloned into pCR blunt II Topo such that SP6 phage promoter was at the 5’ end and restriction site EcoRV was at the 3’ end. The 0.26 kbp insert was sequence-confirmed. Control NB-KO (pNB163) and anti-Rev NB-KO (pNB190) plasmids were kind gifts of Thomas Vercruysse (KU Leuven in Leuven, Belgium). These were already in a CMV expression vector. pME VSV G and HIV-PV were as previously described. pCMV-HIV Gag-RRE was a kind gift of Alan Rein (NCI-Frederick).

### HIV Vectors and cells

Starting third generation HIV transfer vector was pFG12, which was cleaved with Xba1 and Cla1 and ~2.6 kbp CMV-IRES-puro^r^ Xba1+Cla1 fragment from pQCXIP (Clontech) was inserted to construct pHIV-CMV-IRES-puro. This vector was cleaved with BamH1 and Cla1, blunted with Klenow enzyme to insert a 1.6 kb IRES-hygro^r^ cassette to construct pHIV-CMV-IRES-hygro. pHIV-CMV-IRES-puro was cut with EcoR1 and Afe1 to separately insert 1.4 kbp IRES-bleo^r^ and 1.0 kbp IRES-bsd^r^ cassettes to construct pHIV-CMV-IRES-bleo and pHIV-CMV-IRES-blasti, respectively. Each of these HIV vectors has several unique sites just 3’ of the CMV promoter and 5’ of the IRES for transgene insertion; additional details of the construction of each plasmid are available upon request. Rev-NLUC was removed from pRev-NLUC using restriction sites HindIII and EcoRV and blunt-cloned into the unique Age1 site of pHIV-CMV-IRES-puro to make pHIV-CMV-Rev-NLUC-IRES-puro. Similarly, CLUC-Rev was removed from pCLUC-Rev using sites HindIII and EcoRV and blunt-cloned into the Age1 site of pHIV-CMV-IRES-hygro to make HIV-CMV-CLUC-Rev-IRES-hygro. pNB190 were cleaved with HindIII and EcoRV to remove intact the 1.0 kbp NB-KO fusion ORF and blunt-cloned into the unique EcoR1 site of pHIV-CMV-IRES-blasti to construct HIV-CMV-NB190-IRES-blasti.

CMV expression vector encoding Rev-eGFP, initially made by Maria Zapp (UMass Medical Center, Worcester, MA), was obtained from Heinrich Gottlinger (also of UMass). The Rev-eGFP ORF of this plasmid was PCR amplified using primers 5’-GGGCTAGCGCCACCATGGCAGGAAGAAGCGGAGACAGC-3’ and 5’-CCTCTAGATTACTTGTACAGCTCGTCCATGCCGAG-3’, Topo cloned as described above and sequence-confirmed, cut out with EcoR1, and the 1.1 kbp fragment inserted into the unique EcoR1 site of pHIV-CMV-IRES-bleo to create HIV-CMV-ReveGFP-IRES-bleo. To render Rev non-functional in pHIV-PV the plasmid was cleaved with BamH1, Klenow-filled to add four bp and thus frame-shift the Rev ORF, and resealed, with the now-blunted DNA fragment placed back in the original orientation. The resultant plasmid was named pHIV-PVΔRev.

293T cells were maintained in DMEM complete as described. VSV G-pseudotyped particles were produced from 293Ts as described, by calcium phosphate-mediated co-transfection of pME VSV G, pHIV-PV, and the appropriate transfer vector, typically in 10 cm plate format. Selection of transduced cells occurred in the presence of the indicated antibiotic: puromycin (Sigma-Aldrich) at 10 μg/ml, hygromycin (ThermoFisher) at 500 μg/ml, phleomycin (Invivogen) at 200-400 μg/ml, or blasticidin (Invivogen) at 10 μg/ml. In some instances, 293T cells were serially transduced and subsequently maintained in DMEM complete with two or three antibiotics, as indicated. After cellular transduction and antibiotic selection there was no overt cytotoxicity for any of the HIV vectors used here or encoded gene products, and cells were typically passaged twice weekly in DMEM complete plus necessary supplements.

### Luciferase luminometry, immunoblotting, and flow cytometry

Typically plasmids were transiently transfected using the calcium phosphate method into 293Ts in 12- or 24-well format in triplicate. Transfected cells were harvested at 72 h, lysed in buffered 0.1% Triton X-100 solution, and FFLUC measured as described in 96-well format, using a BioTek Synergy™ 2 Microplate Reader. In some cases, RLU values were normalized to wt Rev, set at 100% or 1.0. In order to detect protein products, detergent-treated cell lysates were size-separated on pre-made SDS-PAGE gradient gels (Bio-Rad), transferred to nitrocellulose membranes, and probed with antibody as described. To detect FFLUC-containing fusion proteins, rabbit anti-luciferase antibody (Abcam) was used as the primary at 1:5000, and secondary antibody was goat anti-rabbit conjugated to HRP. To detect HA epitope, anti-HA 12CA5 mouse monoclonal antibody (Sigma-Aldrich) was used at 1:2000 with secondary being anti-mouse-HRP (Sigma-Aldrich) at 1:2000. To detect Rev, anti-Rev mouse monoclonal antibody (1G7 from the NIH AIDS Reagent Program or Rev-4 from Santa Cruz Biotechnology) was used a 1:1000 as primary, and secondary was anti-mouse-HRP at 1:2000. To detect Gag precursor p55, primary antibody was anti-HIV 1984 at 1:2000 (NIH AIDS Reagent Program), with secondary anti-human-HRP (Sigma-Aldrich) at 1:2000.

FACS analysis was used to detect GFP on either the FACSCalibur (Becton Dickinson) or LSRII (BD) instruments, collecting at least 10,000 events. Cell sorting for KO+ cells was performed on an FACS Aria (BD), positively gating on the KO+ cells within the highest fluorescence quartile.

## Acknowledgements

We thank Drs. Maria Zapp, Heinrich Gottlinger, Bryan Cullen, Joel Belasco, Alan Rein, and Thomas Vercruysse for generous reagent gifts. The following reagents were obtained through the NIH AIDS Reagent Program, Division of AIDS, NIAID, NIH: HIV-1 Rev Monoclonal Antibody 1G7 (cat# 7376) from Dr. Anne Marie Szilvay; HIV-1 Neutralizing Serum 1984 from Dr. Luba Vujcic, FDA, Center for Biologics Evaluation and Research. This work was supported by NIH grants DP1 DA036463 and R21 AI136768.

## Notes

### Competing Interest Statement

The authors have declared no competing interest.

## References

1. Marcus JL, Chao CR, Leyden WA, Xu L, Quesenberry CP, Jr., Klein DB, et al. Narrowing the Gap in Life Expectancy Between HIV-Infected and HIV-Uninfected Individuals With Access to Care. J Acquir Immune Defic Syndr. 2016;73(1):39–46. Epub 2016/03/31. doi: 10.1097/qai.0000000000001014. PubMed PMID: 27028501; PubMed Central PMCID: PMCPMC5427712.

2. Siddiqi AE, Hall HI, Hu X, Song R. Population-Based Estimates of Life Expectancy After HIV Diagnosis: United States 2008-2011. J Acquir Immune Defic Syndr. 2016;72(2):230–6. Epub 2016/02/19. doi: 10.1097/qai.0000000000000960. PubMed PMID: 26890283; PubMed Central PMCID: PMCPMC4876430.

3. Cihlar T, Fordyce M. Current status and prospects of HIV treatment. Curr Opin Virol. 2016;18:50–6. Epub 2016/03/30. doi: 10.1016/j.coviro.2016.03.004. PubMed PMID: 27023283.

4. Flepisi BT, Bouic P, Sissolak G, Rosenkranz B. Drug-drug interactions in HIV positive cancer patients. Biomed Pharmacother. 2014;68(5):665–77. Epub 2014/05/28. doi: 10.1016/j.biopha.2014.04.010. PubMed PMID: 24863536.

5. McIlleron H, Abdel-Rahman S, Dave JA, Blockman M, Owen A. Special populations and pharmacogenetic issues in tuberculosis drug development and clinical research. J Infect Dis. 2015;211 Suppl 3(Suppl 3):S115-25. Epub 2015/05/27. doi: 10.1093/infdis/jiu600. PubMed PMID: 26009615; PubMed Central PMCID: PMCPMC4551115.

6. Wyles DL. Regimens for Patients Coinfected with Human Immunodeficiency Virus. Clin Liver Dis. 2015;19(4):689–706, vi-vii. Epub 2015/10/16. doi: 10.1016/j.cld.2015.06.008. PubMed PMID: 26466656.

7. Burgess S, Partovi N, Yoshida EM, Erb SR, Azalgara VM, Hussaini T. Drug Interactions With Direct-Acting Antivirals for Hepatitis C: Implications for HIV and Transplant Patients. Ann Pharmacother. 2015;49(6):674–87. Epub 2015/03/15. doi: 10.1177/1060028015576180. PubMed PMID: 25770114.

8. Meemken L, Hanhoff N, Tseng A, Christensen S, Gillessen A. Drug-Drug Interactions With Antiviral Agents in People Who Inject Drugs Requiring Substitution Therapy. Ann Pharmacother. 2015;49(7):796–807. Epub 2015/04/24. doi: 10.1177/1060028015581848. PubMed PMID: 25902733.

9. Kumar S, Rao PS, Earla R, Kumar A. Drug-drug interactions between anti-retroviral therapies and drugs of abuse in HIV systems. Expert Opin Drug Metab Toxicol. 2015;11(3):343–55. Epub 2014/12/30. doi: 10.1517/17425255.2015.996546. PubMed PMID: 25539046; PubMed Central PMCID: PMCPMC4428551.

10. Vadlapatla RK, Patel M, Paturi DK, Pal D, Mitra AK. Clinically relevant drug-drug interactions between antiretrovirals and antifungals. Expert Opin Drug Metab Toxicol. 2014;10(4):561–80. Epub 2014/02/14. doi: 10.1517/17425255.2014.883379. PubMed PMID: 24521092; PubMed Central PMCID: PMCPMC4516223.

11. Hughes CA, Foisy M, Tseng A. Interactions between antifungal and antiretroviral agents. Expert Opin Drug Saf. 2010;9(5):723–42. Epub 2010/03/30. doi: 10.1517/14740331003752694. PubMed PMID: 20345324.

12. Cooper RD, Wiebe N, Smith N, Keiser P, Naicker S, Tonelli M. Systematic review and meta-analysis: renal safety of tenofovir disoproxil fumarate in HIV-infected patients. Clin Infect Dis. 2010;51(5):496–505. Epub 2010/08/03. doi: 10.1086/655681. PubMed PMID: 20673002.

13. Fortuny C, Deyà-Martínez Á, Chiappini E, Galli L, de Martino M, Noguera-Julian A. Metabolic and renal adverse effects of antiretroviral therapy in HIV-infected children and adolescents. Pediatr Infect Dis J. 2015;34(5 Suppl 1):S36–43. Epub 2015/01/30. doi: 10.1097/inf.0000000000000663. PubMed PMID: 25629891.

14. Lv Z, Chu Y, Wang Y. HIV protease inhibitors: a review of molecular selectivity and toxicity. HIV AIDS (Auckl). 2015;7:95–104. Epub 2015/04/22. doi: 10.2147/hiv.S79956. PubMed PMID: 25897264; PubMed Central PMCID: PMCPMC4396582.

15. Gardner K, Hall PA, Chinnery PF, Payne BA. HIV treatment and associated mitochondrial pathology: review of 25 years of in vitro, animal, and human studies. Toxicol Pathol. 2014;42(5):811–22. Epub 2013/09/27. doi: 10.1177/0192623313503519. PubMed PMID: 24067671.

16. Margolis AM, Heverling H, Pham PA, Stolbach A. A review of the toxicity of HIV medications. J Med Toxicol. 2014;10(1):26–39. Epub 2013/08/22. doi: 10.1007/s13181-013-0325-8. PubMed PMID: 23963694; PubMed Central PMCID: PMCPMC3951641.

17. Abers MS, Shandera WX, Kass JS. Neurological and psychiatric adverse effects of antiretroviral drugs. CNS Drugs. 2014;28(2):131–45. Epub 2013/12/24. doi: 10.1007/s40263-013-0132-4. PubMed PMID: 24362768.

18. Pham QD, Wilson DP, Law MG, Kelleher AD, Zhang L. Global burden of transmitted HIV drug resistance and HIV-exposure categories: a systematic review and meta-analysis. Aids. 2014;28(18):2751–62. Epub 2014/12/11. doi: 10.1097/qad.0000000000000494. PubMed PMID: 25493601.

19. Frentz D, van de Vijver D, Abecasis A, Albert J, Hamouda O, Jørgensen L, et al. Patterns of transmitted HIV drug resistance in Europe vary by risk group. PLoS One. 2014;9(4):e94495. Epub 2014/04/12. doi: 10.1371/journal.pone.0094495. PubMed PMID: 24721998; PubMed Central PMCID: PMCPMC3983178 Editorial Board member. This does not alter the authors’ adherence to PLOS ONE Editorial policies and criteria.

20. Frentz D, Van de Vijver DA, Abecasis AB, Albert J, Hamouda O, Jørgensen LB, et al. Increase in transmitted resistance to non-nucleoside reverse transcriptase inhibitors among newly diagnosed HIV-1 infections in Europe. BMC Infect Dis. 2014;14:407. Epub 2014/07/23. doi: 10.1186/1471-2334-14-407. PubMed PMID: 25047543; PubMed Central PMCID: PMCPMC4223652.

21. Vega Y, Delgado E, Fernández-García A, Cuevas MT, Thomson MM, Montero V, et al. Epidemiological Surveillance of HIV-1 Transmitted Drug Resistance in Spain in 2004-2012: Relevance of Transmission Clusters in the Propagation of Resistance Mutations. PLoS One. 2015;10(5):e0125699. Epub 2015/05/27. doi: 10.1371/journal.pone.0125699. PubMed PMID: 26010948; PubMed Central PMCID: PMCPMC4444345.

22. Global epidemiology of drug resistance after failure of WHO recommended first-line regimens for adult HIV-1 infection: a multicentre retrospective cohort study. Lancet Infect Dis. 2016;16(5):565–75. Epub 2016/02/03. doi: 10.1016/s1473-3099(15)00536-8. PubMed PMID: 26831472; PubMed Central PMCID: PMCPMC4835583.

23. Fogel JM, Hudelson SE, Ou SS, Hart S, Wallis C, Morgado MG, et al. Brief Report: HIV Drug Resistance in Adults Failing Early Antiretroviral Treatment: Results From the HIV Prevention Trials Network 052 Trial. J Acquir Immune Defic Syndr. 2016;72(3):304–9. Epub 2016/02/10. doi: 10.1097/qai.0000000000000951. PubMed PMID: 26859828; PubMed Central PMCID: PMCPMC4911290.

24. Manasa J, Danaviah S, Lessells R, Elshareef M, Tanser F, Wilkinson E, et al. Increasing HIV-1 Drug Resistance Between 2010 and 2012 in Adults Participating in Population-Based HIV Surveillance in Rural KwaZulu-Natal, South Africa. AIDS Res Hum Retroviruses. 2016;32(8):763–9. Epub 2016/03/24. doi: 10.1089/aid.2015.0225. PubMed PMID: 27002368; PubMed Central PMCID: PMCPMC4971422.

25. Malim MH, Böhnlein S, Hauber J, Cullen BR. Functional dissection of the HIV-1 Rev trans-activator--derivation of a trans-dominant repressor of Rev function. Cell. 1989;58(1):205–14. Epub 1989/07/14. doi: 10.1016/0092-8674(89)90416-9. PubMed PMID: 2752419.

26. Malim MH, Hauber J, Le SY, Maizel JV, Cullen BR. The HIV-1 rev trans-activator acts through a structured target sequence to activate nuclear export of unspliced viral mRNA. Nature. 1989;338(6212):254-7. Epub 1989/03/16. doi: 10.1038/338254a0. PubMed PMID: 2784194.

27. Vercruysse T, Daelemans D. HIV-1 Rev multimerization: mechanism and insights. Curr HIV Res. 2013;11(8):623–34. Epub 2014/03/13. doi: 10.2174/1570162x12666140307094603. PubMed PMID: 24606219.

28. Malim MH, Cullen BR. HIV-1 structural gene expression requires the binding of multiple Rev monomers to the viral RRE: implications for HIV-1 latency. Cell. 1991;65(2):241–8. Epub 1991/04/19. doi: 10.1016/0092-8674(91)90158-u. PubMed PMID: 2015625.

29. Madore SJ, Tiley LS, Malim MH, Cullen BR. Sequence requirements for Rev multimerization in vivo. Virology. 1994;202(1):186–94. Epub 1994/07/01. doi: 10.1006/viro.1994.1334. PubMed PMID: 7516596.

30. Szilvay AM, Brokstad KA, Bøe SO, Haukenes G, Kalland KH. Oligomerization of HIV-1 Rev mutants in the cytoplasm and during nuclear import. Virology. 1997;235(1):73–81. Epub 1997/08/18. doi: 10.1006/viro.1997.8671. PubMed PMID: 9300038.

31. Mann DA, Mikaélian I, Zemmel RW, Green SM, Lowe AD, Kimura T, et al. A molecular rheostat. Co-operative rev binding to stem I of the rev-response element modulates human immunodeficiency virus type-1 late gene expression. J Mol Biol. 1994;241(2):193–207. Epub 1994/08/12. doi: 10.1006/jmbi.1994.1488. PubMed PMID: 8057359.

32. Daugherty MD, D’Orso I, Frankel AD. A solution to limited genomic capacity: using adaptable binding surfaces to assemble the functional HIV Rev oligomer on RNA. Mol Cell. 2008;31(6):824–34. Epub 2008/10/17. doi: 10.1016/j.molcel.2008.07.016. PubMed PMID: 18922466; PubMed Central PMCID: PMCPMC2651398.

33. Daly TJ, Cook KS, Gray GS, Maione TE, Rusche JR. Specific binding of HIV-1 recombinant Rev protein to the Rev-responsive element in vitro. Nature. 1989;342(6251):816–9. Epub 1989/12/14. doi: 10.1038/342816a0. PubMed PMID: 2481237.

34. Cook KS, Fisk GJ, Hauber J, Usman N, Daly TJ, Rusche JR. Characterization of HIV-1 REV protein: binding stoichiometry and minimal RNA substrate. Nucleic Acids Res. 1991;19(7):1577–83. Epub 1991/04/11. doi: 10.1093/nar/19.7.1577. PubMed PMID: 2027765; PubMed Central PMCID: PMCPMC333918.

35. Holland SM, Ahmad N, Maitra RK, Wingfield P, Venkatesan S. Human immunodeficiency virus rev protein recognizes a target sequence in rev-responsive element RNA within the context of RNA secondary structure. J Virol. 1990;64(12):5966–75. Epub 1990/12/01. doi: 10.1128/jvi.64.12.5966-5975.1990. PubMed PMID: 2243382; PubMed Central PMCID: PMCPMC248770.

36. Wingfield PT, Stahl SJ, Payton MA, Venkatesan S, Misra M, Steven AC. HIV-1 Rev expressed in recombinant Escherichia coli: purification, polymerization, and conformational properties. Biochemistry. 1991;30(30):7527–34. Epub 1991/07/30. doi: 10.1021/bi00244a023. PubMed PMID: 1854752.

37. Heaphy S, Finch JT, Gait MJ, Karn J, Singh M. Human immunodeficiency virus type 1 regulator of virion expression, rev, forms nucleoprotein filaments after binding to a purine-rich “bubble” located within the rev-responsive region of viral mRNAs. Proc Natl Acad Sci U S A. 1991;88(16):7366–70. Epub 1991/08/15. doi: 10.1073/pnas.88.16.7366. PubMed PMID: 1871141; PubMed Central PMCID: PMCPMC52296.

38. Daugherty MD, Liu B, Frankel AD. Structural basis for cooperative RNA binding and export complex assembly by HIV Rev. Nat Struct Mol Biol. 2010;17(11):1337–42. Epub 2010/10/19. doi: 10.1038/nsmb.1902. PubMed PMID: 20953181; PubMed Central PMCID: PMCPMC2988976.

39. Fang X, Wang J, O’Carroll IP, Mitchell M, Zuo X, Wang Y, et al. An unusual topological structure of the HIV-1 Rev response element. Cell. 2013;155(3):594–605. Epub 2013/11/19. doi: 10.1016/j.cell.2013.10.008. PubMed PMID: 24243017; PubMed Central PMCID: PMCPMC3918456.

40. Fernandes JD, Faust TB, Strauli NB, Smith C, Crosby DC, Nakamura RL, et al. Functional Segregation of Overlapping Genes in HIV. Cell. 2016;167(7):1762–73.e12. Epub 2016/12/17. doi: 10.1016/j.cell.2016.11.031. PubMed PMID: 27984726; PubMed Central PMCID: PMCPMC5287106.

41. Askjaer P, Jensen TH, Nilsson J, Englmeier L, Kjems J. The specificity of the CRM1-Rev nuclear export signal interaction is mediated by RanGTP. J Biol Chem. 1998;273(50):33414–22. Epub 1998/12/05. doi: 10.1074/jbc.273.50.33414. PubMed PMID: 9837918.

42. Neville M, Stutz F, Lee L, Davis LI, Rosbash M. The importin-beta family member Crm1p bridges the interaction between Rev and the nuclear pore complex during nuclear export. Curr Biol. 1997;7(10):767–75. Epub 1997/11/22. doi: 10.1016/s0960-9822(06)00335-6. PubMed PMID: 9368759.

43. Li Y, Bor YC, Misawa Y, Xue Y, Rekosh D, Hammarskjöld ML. An intron with a constitutive transport element is retained in a Tap messenger RNA. Nature. 2006;443(7108):234-7. Epub 2006/09/15. doi: 10.1038/nature05107. PubMed PMID: 16971948.

44. Wang B, Rekosh D, Hammarskjold ML. Evolutionary conservation of a molecular machinery for export and expression of mRNAs with retained introns. Rna. 2015;21(3):426–37. Epub 2015/01/22. doi: 10.1261/rna.048520.114. PubMed PMID: 25605961; PubMed Central PMCID: PMCPMC4338338.

45. Conti E, Franks NP, Brick P. Crystal structure of firefly luciferase throws light on a superfamily of adenylate-forming enzymes. Structure. 1996;4(3):287–98. Epub 1996/03/15. doi: 10.1016/s0969-2126(96)00033-0. PubMed PMID: 8805533.

46. Ozawa T, Kaihara A, Sato M, Tachihara K, Umezawa Y. Split luciferase as an optical probe for detecting protein-protein interactions in mammalian cells based on protein splicing. Anal Chem. 2001;73(11):2516–21. Epub 2001/06/14. doi: 10.1021/ac0013296. PubMed PMID: 11403293.

47. Deluca M. Firefly luciferase. Adv Enzymol Relat Areas Mol Biol. 1976;44:37–68. Epub 1976/01/01. doi: 10.1002/9780470122891.ch2. PubMed PMID: 775940.

48. Li Q, Yoshimura H, Komiya M, Tajiri K, Uesugi M, Hata Y, et al. A robust split-luciferase-based cell fusion screening for discovering myogenesis-promoting molecules. Analyst. 2018;143(14):3472–80. Epub 2018/06/27. doi: 10.1039/c8an00285a. PubMed PMID: 29944152.

49. Sarzotti-Kelsoe M, Bailer RT, Turk E, Lin CL, Bilska M, Greene KM, et al. Optimization and validation of the TZM-bl assay for standardized assessments of neutralizing antibodies against HIV-1. J Immunol Methods. 2014;409:131–46. Epub 2013/12/03. doi: 10.1016/j.jim.2013.11.022. PubMed PMID: 24291345; PubMed Central PMCID: PMCPMC4040342.

50. Luker KE, Smith MC, Luker GD, Gammon ST, Piwnica-Worms H, Piwnica-Worms D. Kinetics of regulated protein-protein interactions revealed with firefly luciferase complementation imaging in cells and living animals. Proc Natl Acad Sci U S A. 2004;101(33):12288–93. Epub 2004/07/31. doi: 10.1073/pnas.0404041101. PubMed PMID: 15284440; PubMed Central PMCID: PMCPMC514471.

51. Yue Y, Coskun AK, Jawanda N, Auer J, Sutton RE. Differential interaction between human and murine Crm1 and lentiviral Rev proteins. Virology. 2018;513:1–10. Epub 2017/10/14. doi: 10.1016/j.virol.2017.09.027. PubMed PMID: 29028476; PubMed Central PMCID: PMCPMC5914484.

52. Jain C, Belasco JG. Structural model for the cooperative assembly of HIV-1 Rev multimers on the RRE as deduced from analysis of assembly-defective mutants. Mol Cell. 2001;7(3):603–14. Epub 2001/07/21. doi: 10.1016/s1097-2765(01)00207-6. PubMed PMID: 11463385.

53. Vercruysse T, Pardon E, Vanstreels E, Steyaert J, Daelemans D. An intrabody based on a llama single-domain antibody targeting the N-terminal alpha-helical multimerization domain of HIV-1 rev prevents viral production. J Biol Chem. 2010;285(28):21768–80. Epub 2010/04/22. doi: 10.1074/jbc.M110.112490. PubMed PMID: 20406803; PubMed Central PMCID: PMCPMC2898381.

54. Dinoso JB, Kim SY, Wiegand AM, Palmer SE, Gange SJ, Cranmer L, et al. Treatment intensification does not reduce residual HIV-1 viremia in patients on highly active antiretroviral therapy. Proc Natl Acad Sci U S A. 2009;106(23):9403–8. Epub 2009/05/28. doi: 10.1073/pnas.0903107106. PubMed PMID: 19470482; PubMed Central PMCID: PMCPMC2685743.

55. Gandhi RT, Zheng L, Bosch RJ, Chan ES, Margolis DM, Read S, et al. The effect of raltegravir intensification on low-level residual viremia in HIV-infected patients on antiretroviral therapy: a randomized controlled trial. PLoS Med. 2010;7(8). Epub 2010/08/17. doi: 10.1371/journal.pmed.1000321. PubMed PMID: 20711481; PubMed Central PMCID: PMCPMC2919424 (consultant); RFS Pharmaceuticals (owns stock options). JJE: Merck, GlaxoSmithKline (consultant and grant support), Bristol-Myers Squibb, Tibotec, Chimerix, Avexa, and Tobira (consultant). RTG: Tibotec (research grant support) and Gilead (grant support). DM: Merck, Bristol-Myers Squibb, and GlaxoSmithKline (consultant and research support); Gilead Sciences (research support and common stock); Chimerix (consultant); Roche, Trimeris, and Tibotec (research support).

56. McMahon D, Jones J, Wiegand A, Gange SJ, Kearney M, Palmer S, et al. Short-course raltegravir intensification does not reduce persistent low-level viremia in patients with HIV-1 suppression during receipt of combination antiretroviral therapy. Clin Infect Dis. 2010;50(6):912–9. Epub 2010/02/17. doi: 10.1086/650749. PubMed PMID: 20156060; PubMed Central PMCID: PMCPMC2897152.

57. Persaud D, Siberry GK, Ahonkhai A, Kajdas J, Monie D, Hutton N, et al. Continued production of drug-sensitive human immunodeficiency virus type 1 in children on combination antiretroviral therapy who have undetectable viral loads. J Virol. 2004;78(2):968–79. Epub 2003/12/25. doi: 10.1128/jvi.78.2.968-979.2004. PubMed PMID: 14694128; PubMed Central PMCID: PMCPMC368798.

58. Shen L, Siliciano RF. Viral reservoirs, residual viremia, and the potential of highly active antiretroviral therapy to eradicate HIV infection. J Allergy Clin Immunol. 2008;122(1):22–8. Epub 2008/07/08. doi: 10.1016/j.jaci.2008.05.033. PubMed PMID: 18602567; PubMed Central PMCID: PMCPMC6812482.

59. Günthard HF, Frost SD, Leigh-Brown AJ, Ignacio CC, Kee K, Perelson AS, et al. Evolution of envelope sequences of human immunodeficiency virus type 1 in cellular reservoirs in the setting of potent antiviral therapy. J Virol. 1999;73(11):9404–12. Epub 1999/10/09. doi: 10.1128/jvi.73.11.9404-9412.1999. PubMed PMID: 10516049; PubMed Central PMCID: PMCPMC112975.

60. Kearney MF, Spindler J, Shao W, Yu S, Anderson EM, O’Shea A, et al. Lack of detectable HIV-1 molecular evolution during suppressive antiretroviral therapy. PLoS Pathog. 2014;10(3):e1004010. Epub 2014/03/22. doi: 10.1371/journal.ppat.1004010. PubMed PMID: 24651464; PubMed Central PMCID: PMCPMC3961343 Pharmaceuticals. This does not alter our adherence to all PLoS Pathogens policies on sharing data and materials.

61. Evering TH, Mehandru S, Racz P, Tenner-Racz K, Poles MA, Figueroa A, et al. Absence of HIV-1 evolution in the gut-associated lymphoid tissue from patients on combination antiviral therapy initiated during primary infection. PLoS Pathog. 2012;8(2):e1002506. Epub 2012/02/10. doi: 10.1371/journal.ppat.1002506. PubMed PMID: 22319447; PubMed Central PMCID: PMCPMC3271083 Healthcare/GlaxoSmithKline. He is also a paid speaker for Gilead and Tibotec and receives Research Support from Gilead. The remaining authors have declared that no competing interests exist.

62. Lorenzo-Redondo R, Fryer HR, Bedford T, Kim EY, Archer J, Pond SLK, et al. Persistent HIV-1 replication maintains the tissue reservoir during therapy. Nature. 2016;530(7588):51-6. Epub 2016/01/28. doi: 10.1038/nature16933. PubMed PMID: 26814962; PubMed Central PMCID: PMCPMC4865637.

63. Fletcher CV, Staskus K, Wietgrefe SW, Rothenberger M, Reilly C, Chipman JG, et al. Persistent HIV-1 replication is associated with lower antiretroviral drug concentrations in lymphatic tissues. Proc Natl Acad Sci U S A. 2014;111(6):2307–12. Epub 2014/01/29. doi: 10.1073/pnas.1318249111. PubMed PMID: 24469825; PubMed Central PMCID: PMCPMC3926074.

64. Petta I, Lievens S, Libert C, Tavernier J, De Bosscher K. Modulation of Protein-Protein Interactions for the Development of Novel Therapeutics. Mol Ther. 2016;24(4):707–18. Epub 2015/12/18. doi: 10.1038/mt.2015.214. PubMed PMID: 26675501; PubMed Central PMCID: PMCPMC4886928.

65. Hong B, van den Heuvel AP, Prabhu VV, Zhang S, El-Deiry WS. Targeting tumor suppressor p53 for cancer therapy: strategies, challenges and opportunities. Curr Drug Targets. 2014;15(1):80–9. Epub 2014/01/07. doi: 10.2174/1389450114666140106101412. PubMed PMID: 24387333.

66. Estrada-Ortiz N, Neochoritis CG, Domling A. How To Design a Successful p53-MDM2/X Interaction Inhibitor: A Thorough Overview Based on Crystal Structures. ChemMedChem. 2016;11(8):757–72. Epub 2015/12/18. doi: 10.1002/cmdc.201500487. PubMed PMID: 26676832; PubMed Central PMCID: PMCPMC4838565.

67. Zhang B, Golding BT, Hardcastle IR. Small-molecule MDM2-p53 inhibitors: recent advances. Future Med Chem. 2015;7(5):631–45. Epub 2015/04/30. doi: 10.4155/fmc.15.13. PubMed PMID: 25921402.

68. Zhao Y, Aguilar A, Bernard D, Wang S. Small-molecule inhibitors of the MDM2-p53 protein-protein interaction (MDM2 Inhibitors) in clinical trials for cancer treatment. J Med Chem. 2015;58(3):1038–52. Epub 2014/11/15. doi: 10.1021/jm501092z. PubMed PMID: 25396320; PubMed Central PMCID: PMCPMC4329994.

